# Medial entorhinal-hippocampal desynchronization parallels the emergence of memory impairment in a mouse model of Alzheimer’s disease pathology

**DOI:** 10.1101/2025.01.15.633171

**Authors:** Lauren M. Vetere, Angelina M. Galas, Nick Vaughan, Yu Feng, Zoé Christenson Wick, Paul A. Philipsberg, Olga Liobimova, Antonio Fernandez-Ruiz, Denise J. Cai, Tristan Shuman

## Abstract

Alzheimer’s disease (AD) is a neurodegenerative disease characterized by progressive impairments in episodic and spatial memory, as well as circuit and network-level dysfunction. While functional impairments in medial entorhinal cortex (MEC) and hippocampus (HPC) have been observed in patients and rodent models of AD, it remains unclear how communication between these regions breaks down in disease, and what specific physiological changes are associated with the onset of memory impairment. We used silicon probes to simultaneously record neural activity in MEC and hippocampus before or after the onset of spatial memory impairment in the 3xTg mouse model of AD pathology. We found that reduced hippocampal theta power, reduced MEC-CA1 theta coherence, and altered phase locking of MEC and hippocampal neurons all coincided with the emergence of spatial memory impairment in 3xTg mice. Together, these findings indicate that disrupted temporal coordination of neural activity in the MEC-hippocampal system parallels the emergence of memory impairment in a model of AD pathology.

GRAPHICAL ABSTRACT

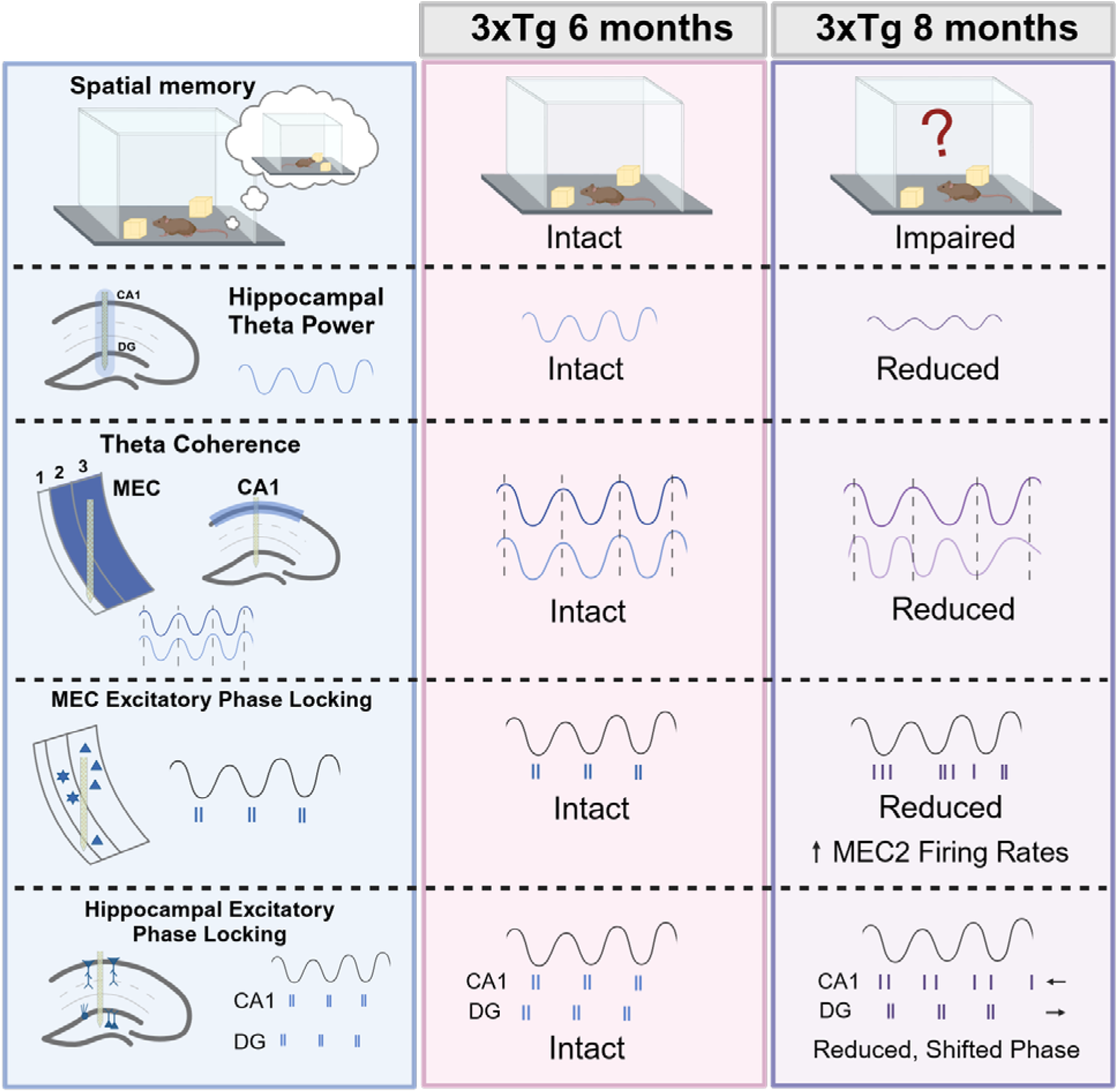

**Highlights:**

- Spatial memory impairments in 3xTg mice emerge between 6 and 8 months of age
- Memory deficits emerge alongside reduced HPC theta power and reduced CA1-MEC theta coherence
- Memory deficits emerge alongside disrupted timing of MEC and HPC neural activity
- Interneuron loss in MEC and interneuron desynchronization in HPC occur prior to memory impairment

## Introduction

Alzheimer’s disease (AD) is a neurodegenerative disease characterized by progressive age-related cognitive impairments and circuit dysfunction. Recent efforts to find new treatments have largely focused on removing pathological proteins, such as amyloid-beta (Aβ), from the brain, but such therapies have shown limited ability to slow cognitive decline^1,2^. An alternative approach that addresses underlying changes in the neural circuits that support memory may be necessary to preserve or restore brain function. In order to design effective interventions, we must first identify specific alterations in circuit function that coincide with the onset of memory impairment and are thus potential drivers of cognitive decline.

Deficits in spatial memory and navigation are among the earliest and most common manifestations of cognitive impairment in AD^3,4^. This suggests that the neural circuits controlling spatial memory are particularly vulnerable to the pathological processes driving this disease. Indeed, the hippocampus and medial entorhinal cortex (MEC) are both critical for spatial memory^5–7^ and exhibit extensive dysfunction in AD patients and rodent models. There is substantial work demonstrating that AD pathology disrupts hippocampal excitability, plasticity, synaptic function, and place cell coding in rodents^8–15^. In addition, upstream inputs to hippocampus from the entorhinal cortex are similarly vulnerable to hyperexcitability, early accumulation of pathology, and cell loss^16–23^. In MEC in particular, recent work in animal models has found early deficits in spatial coding of grid cells^14,24^. Together, this work suggests that disrupted function within the hippocampus-MEC circuit is likely to contribute to the spatial memory impairments observed in AD patients and animal models of AD pathology.

Successful spatial memory and spatial coding depend upon precise timing and synchronization of neural activity within and across the hippocampus and MEC^7,25,26^. Synchrony of neural oscillations and precise timing of neuronal firing support plasticity and facilitate information flow within this circuitry^27–32^. In AD patients and animal models, extensive network changes disrupt synchrony within these brain regions, manifesting as decreased oscillatory power, epileptiform activity, and even seizures^14,33–41^. While network dysfunction is a critical component of this disease, it remains unclear when these changes develop across disease progression, how synchrony is disrupted across regions, and how these changes relate to cognitive decline.

Although dysfunction in entorhinal cortex and hippocampus is well established in AD, the precise cell types and subcircuits that are most impaired within the MEC and hippocampus are not fully understood. Inputs from MEC to hippocampus are made up of two primary projections^42,43^: the “direct pathway” from MEC layer 3 (MEC3) principal cells to CA1^44^, and the “indirect pathway,” which consists of MEC layer 2 (MEC2) stellate cells that project to the dentate gyrus (DG)^45^. These projections provide the major source of spatially tuned input to the hippocampus, including grid, border, head direction, and speed cells^46–50^. Determining which specific circuits are altered across disease progression is key to understanding how memory deficits arise in AD.

To assess the impact of Alzheimer’s disease pathology on hippocampal-entorhinal circuits, we used 3xTg mice, which express APP_SWE_, PS1_M146V_, and Tau_P301L_ mutations. Together, these mutations drive accumulation of Aβ and hyperphosphorylated tau throughout the brain, as well as progressive memory impairments with age^51^. To examine whether network synchrony and spike timing in the hippocampus-MEC circuit are disrupted in these animals, we performed simultaneous *in vivo* electrophysiology recordings in MEC and hippocampus at time points immediately before or after the onset of spatial memory impairments. This approach allowed us to capture network activity and single unit firing in MEC and hippocampus at the onset of spatial memory deficits. We found several critical disruptions in MEC-hippocampal synchronization that emerged alongside memory impairments, including reduced hippocampal theta power, reduced theta coherence between MEC and hippocampus, hyperactivity in MEC2 single units, and disorganized single unit spike timing in MEC and hippocampus. This loss of synchrony and coordinated neuronal firing in MEC and hippocampus is likely to impair spatial memory functions that rely on precise timing in these circuits^7,25,26^. Together, these findings demonstrate impaired MEC-hippocampal communication in 3xTg mice with a time course that parallels the onset of spatial memory impairments.

## Results

### Spatial memory impairments emerge between 6 and 8 months of age in 3xTg mice

We first set out to characterize the time course of spatial memory deficits in 3xTg mice. To assess spatial memory, we utilized a Novel Object Location task, which has been found to be dependent on both the hippocampus and MEC^6,52–54^. Following two sessions of habituation to a square (1 ft x 1 ft) chamber with spatial cues on each wall, animals were placed back in the chamber with two identical objects and allowed to explore the objects for a total of 30 seconds of object exploration. They were then returned to their home cage for 4 hours before being placed back in the chamber with one of the objects moved to a new location (Fig. 1A). We measured how much time each animal spent exploring the moved object as an index of spatial memory. At 6-months of age, there was no difference between 3xTg mice and WT controls in task performance. However, at 8-months-old, 3xTg mice performed significantly worse than their age-matched WT counterparts (Fig. 1B). Notably, both time points are prior to the emergence of aggregated Aβ pathology. Little to no amyloid plaques were detected in 6 or 8-month-old animals, but plaques developed in hippocampus and subiculum by 15 months of age (Fig. 1C-D). This timing aligns with recent phenotypic characterization of this mouse line which showed plaque formation in the subiculum/CA1 at 12 months, though soluble APP and Aβ have been detected by 6-months of age, prior to plaque accumulation^55–58^. Similarly, no neurofibrillary tangles are present at these time points, though phosphorylated tau in subiculum and CA1 has been detected by 6-months of age^55,59–61^. Overall, we found that spatial memory impairments emerged between 6 and 8 months of age in 3xTg mice, prior to Aβ plaque or neurofibrillary tangle formation.

**Figure 1:**
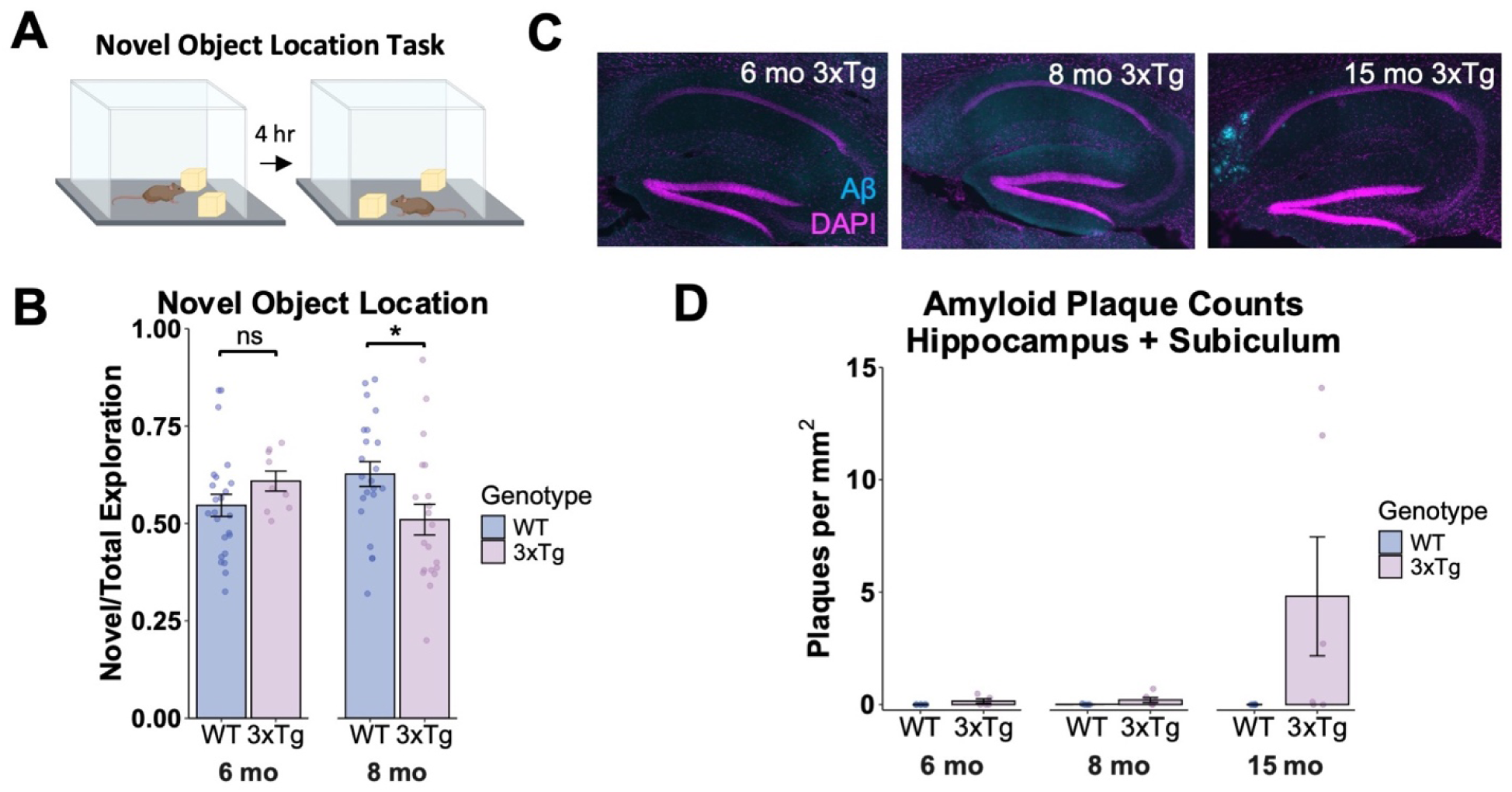
Spatial memory impairment emerges at 8-months old in 3xTg mice. **A.** Schematic depicting the novel object location task. **B.** Performance on the novel object location task. At 6-months, there was no difference in performance between genotypes. However, at 8-months old, 3xTg mice performed significantly worse that age-matched WT controls (Two-way ANOVA, Age x Genotype Interaction: F(1,71) = 5.951, p = 0.017, post hoc tests with Holm correction: WT 6 mo vs 3xTg 6 mo, p= 0.284; WT 8 mo vs 3xTg 8 mo, p =0.025). Sample sizes: WT 6 mo (N= 24, 13 Male and 11 Female), 3xTg 6 mo (N= 9, 4M and 5F), WT 8 mo (N= 22, 5M and 17F), 3xTg 8 mo (N= 20, 8M and 12F). **C.** Representative images from Aβ immunohistochemistry showing no plaques at 6 or 8-months old in 3xTg mice and plaques emerging in subiculum and CA1 by 15 months old. **D.** Quantification of Aβ immunohistochemistry. Sample sizes: WT 6 mo (N = 6, 4M and 2F), 3xTg 6 mo (N = 5, 2M and 3F), WT 8 mo (N = 4, 2M and 2F), 3xTg 8 mo (N = 6, 3M and 3F), WT 15 mo (N = 5, 2M and 3F), 3xTg 15 mo (N = 6, 3M and 3F). Error bars represent s.e.m, *p<0.05. Additional details of all statistical tests can be found in Supplemental Table 1.

Therefore, these time points are ideal to examine how spatial memory circuits are disrupted across the development of memory impairments.

### Reduced hippocampal theta power emerges between 6 and 8 months of age in 3xTg mice

To determine the specific changes in MEC-hippocampal circuit physiology that emerge across the progression of memory deficits, we performed *in vivo* silicon probe recordings in 3xTg and WT mice at 6 or 8 months of age (Fig. 2). We first trained head-fixed mice to run on a virtual reality (VR) linear track to receive water rewards and then performed simultaneous acute electrophysiology in MEC and hippocampus using two 256-channel silicon probes (see Fig. S1 for histological verification of probe locations).

**Figure 2:**
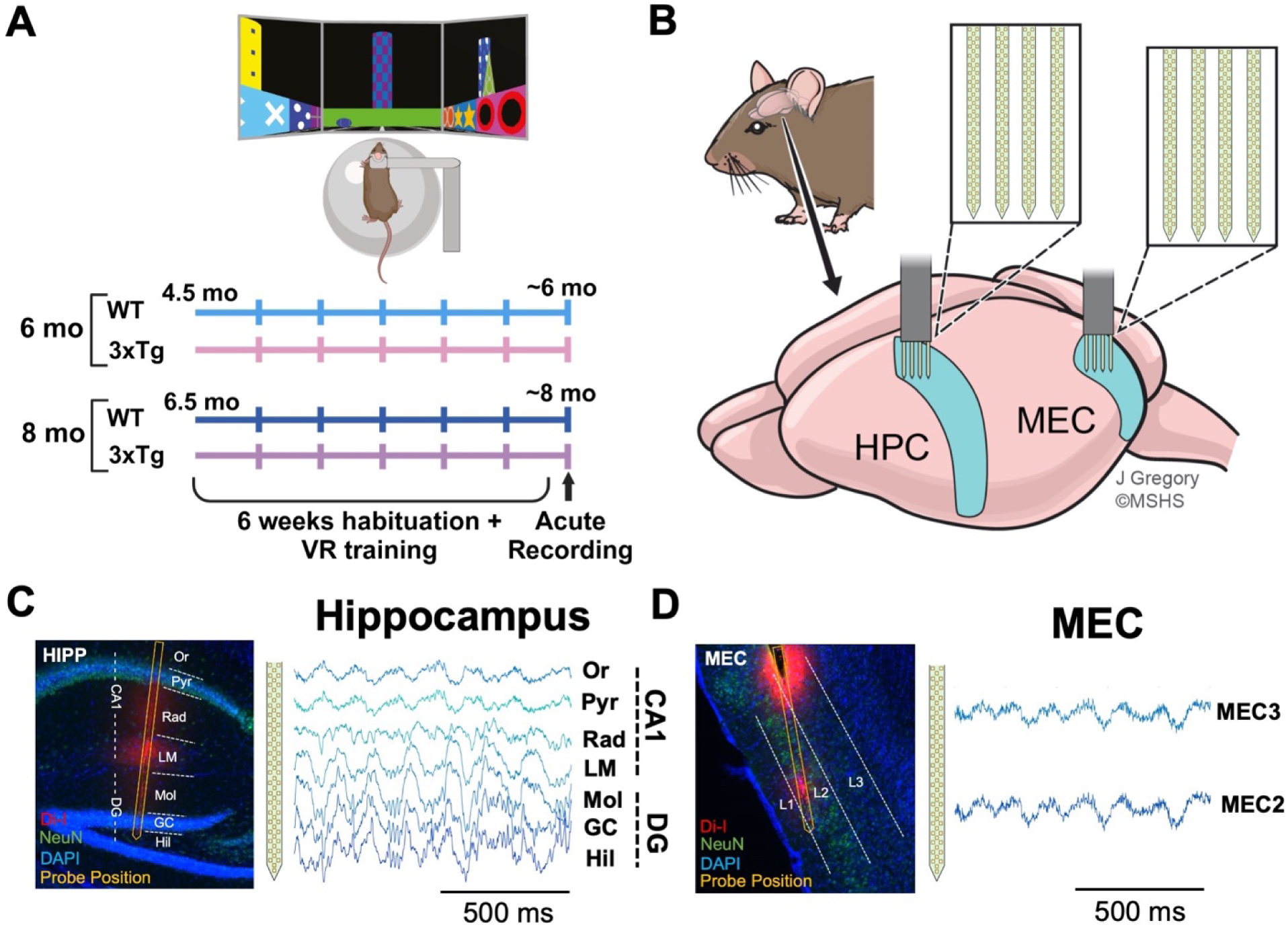
Simultaneous *in vivo* electrophysiology in MEC and hippocampus of WT and 3xTg mice using silicon probes. **A.** Schematic depicting the timeline of electrophysiology experiments. Animals were trained to run while headfixed in a virtual reality environment to receive water rewards. Once an animal was well trained, we inserted silicon probes into both MEC and Hippocampus (HPC) and performed an acute recording while the mouse ran on the virtual track. Animals were recorded at either 6-or 8-months old. **B.** Schematic showing targeted probe locations in MEC and CA1 (4 shanks per probe, 64 channels per shank, 512 channels total across two probes). Shanks span medial to lateral. **C. Left:** Example histology showing one of four shanks in a sagittal slice of hippocampus. Layer labels: Or = stratum oriens, Pyr = stratum pyramidale, Rad = stratum radiatum, LM = stratum lacunosum moleculare, Mol = molecular layer, GC = granule cell layer, Hil = hilus. Probes were painted with Di-I prior to recording to facilitate later histological verification of probe locations. **Right:** Example local field potentials (LFPs) recorded from each layer of the hippocampus, 1 second in length. **D. Left:** Example histology showing one of four shanks in MEC, spanning MEC2 and MEC3 (L1= Layer 1, L2 = Layer 2, L3 = Layer 3). **Right:** Example LFP signals from MEC2 and MEC3, 1 second in length.

We first sought to quantify changes in hippocampal network activity by examining theta oscillations, which are prominent in the temporal lobe and have been associated with successful learning and memory^62–67^. Broadly, theta oscillations are rhythmic extracellular voltage fluctuations that reflect the summation of synaptic activity near the recording electrode. The power, or amplitude, of hippocampal theta oscillations depends largely on transmembrane currents elicited by long-range inputs from the MEC^63,68^. Notably, theta power is influenced by running speed, and thus can be confounded by potential differences in running behavior between groups. While we found no difference is locomotion between WT and 3xTg mice, we did find that older mice of both genotypes ran slower on the virtual linear track (Fig. S2A). Thus, to allow comparisons across both genotype and age, we subsampled the data for periods where running speeds were comparable across groups and used this subsampled data for all analyses (Fig. S2C).

At 6 months of age, prior to the onset of spatial memory impairments, we found no significant differences in hippocampal theta power, or amplitude, between 3xTg and WT mice (Fig. 3A-C). However, at 8 months of age, 3xTg mice had reduced theta power across all CA1 and DG layers (Fig. 3D). Reduced hippocampal theta power is consistent with a loss of the entorhinal inputs that target distal dendrites in CA1 and DG and elicit local theta modulated currents there^63,68^. However, changes in theta power may also reflect changes in volume conducted theta from nearby brain regions. To more clearly define the source of reduced theta power, we performed current source density analysis (CSD) to isolate local current sources and sinks generated along the hippocampal laminar profile^69,70^. Again, at 6 months of age we found no changes in CSD magnitude between genotypes (Fig. 3B-C). In 8-month-old 3xTg mice, however, CSD magnitude was reduced, most prominently in layers that receive input from entorhinal cortex (Fig. 3E-F). In MEC, we found no significant differences in theta power across genotypes at either time point (Fig. S3C-D). Together, these findings suggest that reduced hippocampal theta power, likely driven by disrupted entorhinal input into hippocampus, emerges between 6 and 8 months of age in 3xTg mice, coinciding with the onset of spatial memory impairment in this model.

**Figure 3:**
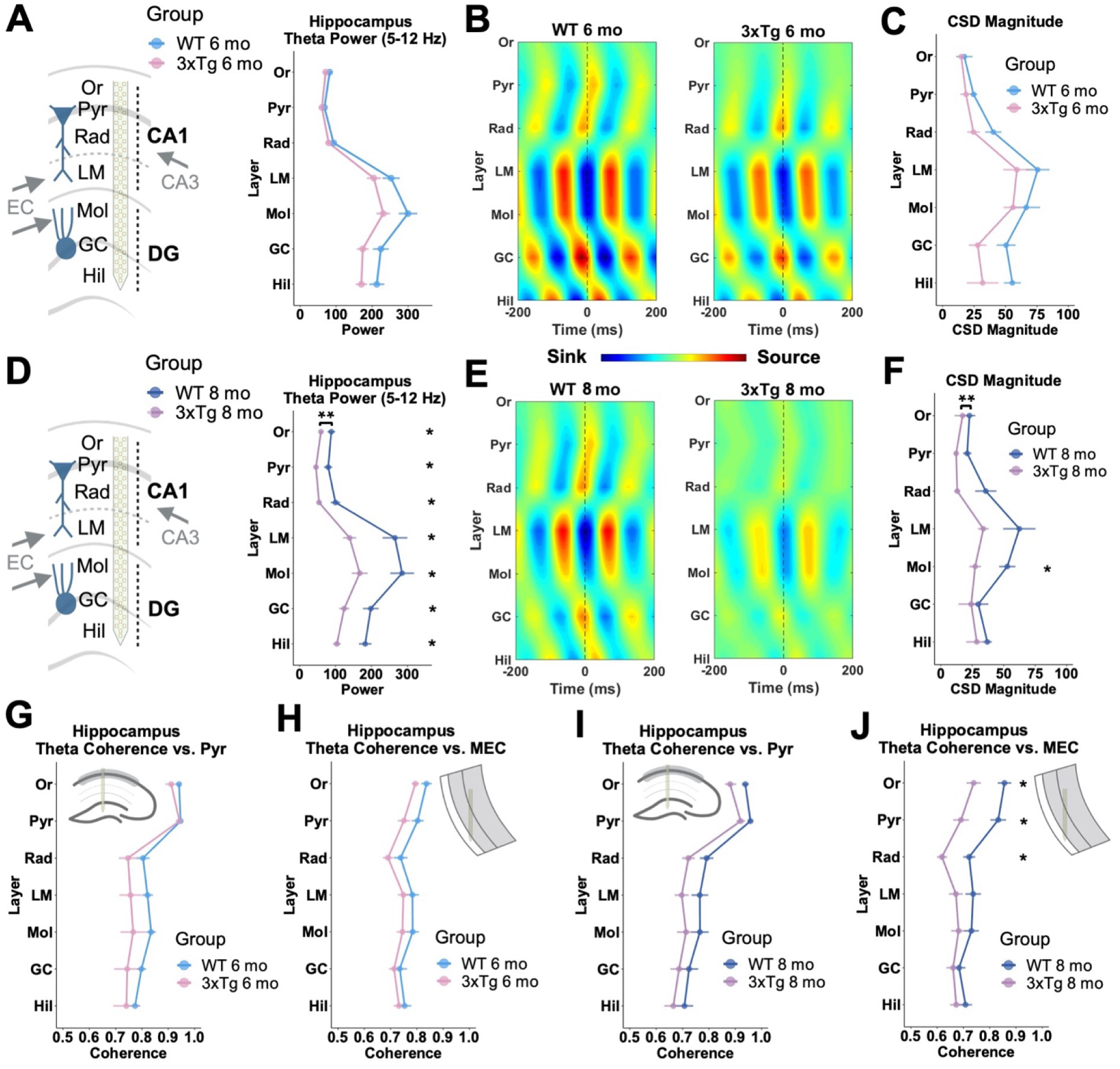
Reduced hippocampal theta power and MEC-CA1 theta coherence in 8-month-old 3xTg mice. **A. Left:** Schematic showing silicon probe location spanning from oriens to hilus in the hippocampus. MEC inputs come in to CA1 lacunosum moleculare (LM) and DG molecular layer (Mol). CA3 inputs come in to CA1 radiatum (Rad). **Right:** Theta power across all layers of hippocampus in WT and 3xTg mice at 6-months old. There were no significant changes in theta power in any hippocampal layer in 6-month-old 3xTg mice compared to age-matched WT controls. (Two-way repeated measures ANOVA, Genotype x Layer Interaction F(6, 90) = 2.26, p = 0.044, posthocs corrected with Holm method, all layers p>0.1). **B.** Theta current source density (CSD) averaged across theta cycles, depicting current sources and sinks in WT 6 mo and 3xTg 6 mo mice. **C.** Maximum CSD magnitude by layer in WT 6 mo and 3xTg 6 mo animals. No significant differences in CSD magnitude across genotypes. **D.** Theta power across hippocampal layers in 8-month-old WT and 3xTg mice. Theta power was reduced throughout the hippocampus in 8-month-old 3xTg mice compared to age-matched controls (Two-way RM ANOVA, Main Effect of Genotype: F(1,15) = 8.67, p <0.05, and Genotype x Layer Interaction: F(6,90) = 5.25, p < 0.001, posthocs corrected with Holm method, p< 0.05 for all layers). **E.** Theta CSD averaged across theta cycles, depicting current sources and sinks in WT 6 mo and 3xTg 8 mo mice. **F.** Maximum CSD magnitude by layer in 8-month-old WT and 3xTg animals. Reduced CSD magnitude in 3xTg mice (Mixed Effects Model, Main Effect of Genotype F(1,15) = 4.91, p = 0.043, posthocs corrected with Holm-Sidak method, Mol: p = 0.042, other layers p> 0.05). **G.** Theta coherence within hippocampus in 6-month-old 3xTg and WT mice. Inset indicates that coherence is calculated between theta in each hippocampal layer and theta in CA1 pyramidal layer. No changes in coherence between genotypes. **H.** Theta coherence between hippocampus and MEC in 6-month-old 3xTg and WT mice. Inset indicates that each row depicts theta coherence between theta in the given hippocampal layer and theta in MEC. No changes between genotypes. **I.** Same as G for 8-month-old groups. No changes in intra-hippocampal coherence between genotypes. **J.** Same as H for 8-month-old groups. Decreased theta coherence between MEC and CA1 (Two-way RM ANOVA, Genotype x Layer interaction F(6,84) = 3.02, p = 0.010, posthocs corrected with Holm method, Rad p =0.027, Pyr p = 0.03, Or p=0.038, other layers p>0.1). N = 9 WT 6mo (5M, 4 F), 8 3xTg 6 mo (3M, 5F), 10 WT 8 mo (5M, 5F), 7 3xTg 8 mo (3M, 4F). One WT 8mo M mouse and one 3xTg 6 M mouse were excluded from H and I due to poor probe targeting in MEC. Two WT 6 mo and one 3xTg 8 mo animal are missing data from oriens in CSD analysis due to limited channels recorded in this layer. Error bars represent s.e.m, *p<0.05, **p<0.01. Additional details of all statistical tests can be found in Supplemental Table 1.

Oscillations at other frequencies are also relevant for cognition and have been implicated in various mouse models of AD pathology^37,71–73^. For this reason, we also examined hippocampal fast gamma (90-130 Hz) and slow gamma (30-50 Hz). Fast gamma is associated with MEC-hippocampus communication, while slow gamma has been linked to CA3◊CA1 and lateral entorhinal cortex◊DG projections^7,74–77^. We found decreased fast gamma power in 8-month-old 3xTg mice, with the most pronounced differences localized in granule cell layer and hilus (Fig. S4C), suggesting that MEC inputs to DG may be altered in these mice. Oscillations in the slow gamma band were relatively unaffected (Fig. S4B,D). Together, these findings are consistent with a disruption of MEC inputs to hippocampus across the development of memory impairments in 3xTg mice.

### Reduced MEC-CA1 theta coherence emerges between 6 and 8 months of age in 3xTg mice

We next sought to assess whether theta oscillations between MEC and hippocampus become desynchronized across the development of memory impairments in 3xTg mice. Theta synchrony between hippocampus and cortical regions is thought to support learning and memory by organizing spiking across connected brain regions and promoting plasticity^30^. To assess synchrony, we calculated the phase coherence (how consistent the phase lag is between two recording sites) of theta oscillations across hippocampus and MEC. In 6-month-old animals, we found no changes in theta coherence between MEC and hippocampus or within hippocampus in 3xTg mice (Fig. 3G-H). However, at 8 months of age, 3xTg mice had reduced theta coherence between MEC and CA1 compared to their age-matched WT controls (Fig. 3J). In particular, theta coherence between MEC and the stratum oriens, pyramidale, and radiatum layers of CA1 was decreased, while coherence between MEC and DG was unaltered. Meanwhile theta coherence within the hippocampus itself was relatively spared (Fig. 3G and 3I), as was coherence within MEC (Fig. S3B). Taken together, these findings suggest a specific deficit in synchrony between MEC and CA1 in 8-month-old 3xTg mice that aligns with the onset of memory impairment.

To investigate whether changes in theta frequency could contribute to deficits in coherence, we calculated the power spectrum density (PSD) within the theta range (5-12Hz) for each animal in MEC and each CA1 sublayer and found the frequency with the highest power within that range. We found a decrease in peak theta frequency in 8-month-old 3xTg mice compared to age-matched WTs that was present in MEC and CA1 radiatum (Fig. S5). Slowing of the theta rhythm in MEC could contribute to reduced entorhinal-hippocampal synchrony, however given that coherence changes were sublayer specific (Fig. 3J), this slowing is unlikely to be the sole driver of theta coherence changes in this circuit. Overall, our analysis of theta oscillations found that reduced hippocampal theta power, CSD magnitude, and MEC-CA1 coherence all emerged in 3xTg mice at 8-months of age, coincident with the onset of spatial memory impairments on the novel object location task. Together, this suggests a progressive breakdown in coordination between MEC and hippocampus in 3xTg mice that may contribute to spatial memory impairments, but it remains unclear which neuronal populations could underlie these effects.

### Loss of MEC3 spike timing precision in 8-month-old 3xTg mice

Reduced hippocampal theta power and loss of coherence between MEC and CA1 may reflect a loss of properly timed inputs to hippocampus, as rhythmic inputs from MEC are critical for hippocampal theta oscillations^63,68^.Thus, we hypothesized that firing patterns of neurons in upstream MEC might be disrupted in 8-month-old 3xTg mice. To assess this, we isolated single units recorded in MEC and separated them into putative excitatory and inhibitory units based on their firing properties and waveform shapes (Fig. S6, see Methods for further details)^78^. We examined the synchrony of individual neurons relative to ongoing network activity by assessing theta phase locking. Specific cell types within the hippocampus and entorhinal cortex tend to fire at particular theta phases, and this coordination is thought to promote plasticity and information flow within this circuitry^78,79^. Here, we quantified the strength of phase locking by measuring the mean resultant vector, or R-value, for each unit. Cells with lower R values have a weaker preference for their preferred firing phase (see additional visualizations illustrating this in Fig. S7). Furthermore, by examining phase locking relative to both local (MEC) or long-range (HPC) theta oscillations, we can gain insight into whether desynchronization emerges locally or specifically between MEC and hippocampus.

Given the specific deficit we found in coherence between MEC and CA1, we first investigated the firing properties of MEC3 excitatory neurons, which are known to directly project to CA1 (Fig. 4A-D). We first asked whether overall activity levels were altered and found no significant differences in firing rates across groups (Fig. 4B). We then examined phase locking to local MEC theta oscillations as a metric of local synchrony and function within MEC (Fig. 4C). Reduced phase locking of MEC3 neurons with MEC theta would indicate reduced coordination of MEC3 spiking with upstream inputs to MEC and with overall local MEC activity. We found no differences in phase locking to local MEC theta between 3xTg and WT groups at 6 months of age. However, excitatory MEC3 neurons in 3xTg mice had reduced phase locking strength between 6 and 8 months of age, while WT neurons did not change across these time points. A direct comparison between 3xTg and WT excitatory MEC3 phase locking at 8 months of age approached significance but did not reach criterion after adjusting for multiple comparisons (Supplemental Table 1). Thus, MEC3 phase locking to local theta weakened with age in 3xTg mice, but not in WT mice.

**Figure 4:**
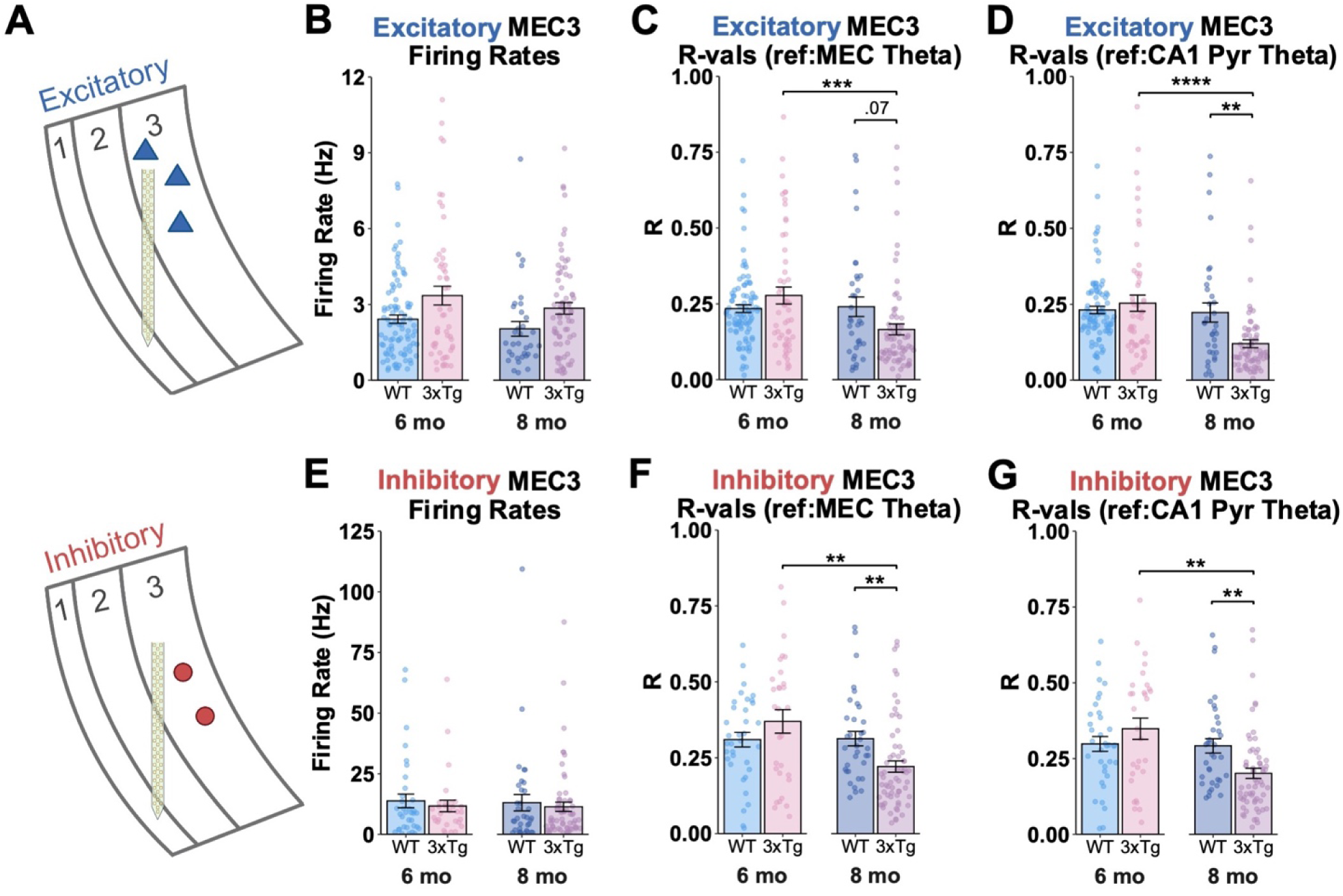
Theta phase locking of MEC3 units is reduced with age in 3xTg mice. **A.** Schematics showing probe location and cell types in MEC3. Excitatory principal cells in MEC3 are shown in the top row in blue, while interneurons are shown in the bottom row in red. **B.** Firing rates of MEC3 putative excitatory neurons. No significant group differences. **C.** Phase locking strength of excitatory MEC3 units to theta oscillations recorded in MEC. R-values were reduced with age in 3xTg mice (3xTg 6 mo vs. 3xTg 8 mo Mann-Whitney (MW) test, adjusted p<0.001, WT 8 mo vs. 8 mo 3xTg adjusted p = 0.07). **D.** Phase locking strength of excitatory MEC3 units to theta recorded in CA1 pyramidal layer. R-values were reduced with age in 3xTg mice. Phase locking to CA1 was decreased in 8-month-old 3xTg animals relative to WT controls. (3xTg 6 mo vs. 3xTg 8 mo MW, adjusted p<0.0001, WT 8 mo vs. 3xTg 8 mo adjusted p = 0.003). **E.** MEC3 inhibitory firing rates. No change in firing rates of inhibitory MEC3 units across groups. **F.** Phase locking strength of inhibitory MEC3 units to theta oscillations recorded in MEC. R-values were reduced with age in 3xTg mice and reduced in 8-month-old 3xTg mice compared to age-matched WT controls (3xTg 6 mo vs. 3xTg 8 mo MW, adjusted p = 0.002, WT 8 mo vs. 3xTg 8 mo adjusted p = 0.005). **G.** Phase locking strength of MEC3 inhibitory units to theta recorded in CA1 pyramidal layer. MEC3 inhibitory units in 8-month-old 3xTg mice exhibit reduced phase locking to CA1 theta (3xTg 6 mo vs. 3xTg 8 mo MW, adjusted p = 0.002, WT 8 mo vs. 3xTg 8 mo adjusted p = 0.002). **Excitatory Units:** WT 6 mo (n = 92), 3xTg 6 mo (n =54), WT 8 mo cells (n = 34), 3xTg 8 mo (n = 71). **Inhibitory Units:** WT 6 mo (n = 35), 3xTg 6 mo (n = 30), WT 8 mo cells (n = 63), 3xTg 8 mo (n = 63). Extended information about # of cells per animal can be found in Supplemental Table 1. Error bars show s.e.m. *p<0.05, **p<0.01, ***p<0.001, ****p<0.0001. Because single unit data did not meet ANOVA assumptions for normality and equal variances, Mann-Whitney tests were used for all comparisons (see Methods and Supplemental Table 1 for more details).

We next examined phase locking of MEC3 excitatory neurons to downstream CA1 theta, in order to assess synchrony between the spiking activity of MEC neurons and neural activity in hippocampus (Fig. 4D). We found no differences in phase locking at 6 months of age, but significant deficits by 8 months of age. Excitatory MEC3 phase locking in 8-month-old 3xTg animals was reduced compared to both age-matched WT controls and younger 6-month-old 3xTg mice. Together, these data demonstrate that MEC3 excitatory neurons have reduced theta phase locking to both local and downstream theta that emerges between 6 and 8 months of age in 3xTg mice (Fig 4C-D). This suggests a loss of synchronization within MEC as well as between MEC3 neurons and their downstream targets in hippocampus.

To gain a more complete picture of MEC3 microcircuitry, we also examined firing properties of MEC3 interneurons, which provide inhibition to local excitatory populations. In inhibitory MEC3 neurons, we again found no changes in firing rates (Fig 4E), and a similar pattern of decreased theta phase locking in 3xTg neurons at 8 months of age when compared to neurons from 6-month-old 3xTg mice or neurons from age-matched WT mice (Fig. 4F-G). These interneuron phase locking deficits further highlight a progressive vulnerability of MEC3 neurons with age in 3xTg mice. Overall, these results point to a desynchronization between MEC3 spike timing and both local and hippocampal theta oscillations that coincides with the onset of spatial memory impairment in 3xTg mice.

### Hyperexcitability and disrupted theta phase locking in MEC2 of 3xTg mice

MEC2 also sends critical projections to hippocampus and contains neuronal populations that have been implicated in AD^16,18–20,22,45,80^. Thus, we next focused our analysis on MEC2 excitatory neurons (Fig. 5A-D), which include a mix of both stellate cells and pyramidal cells with distinct molecular and anatomical features (however, these cell types cannot be distinguished within our recordings). MEC2 stellate cells make up the major projection to DG^45^ and have been shown to be vulnerable to pathology in early AD^16,19,20,81^. We have previously found that MEC2 stellate cells are uniquely hyperexcitable in 10-month-old 3xTg mice *in vitro*^22^, but their *in vivo* firing patterns have not been investigated. To address this, we first asked whether MEC2 excitatory neurons had increased activity during spatial exploration *in vivo*. We found increased firing rates in excitatory MEC2 cells in 8-month-old 3xTg mice (Fig. 5B, but no changes in firing rate at 6-months of age, indicating that MEC2 excitatory neuron hyperactivity emerged across the progression of memory impairments in 3xTg mice.

**Figure 5:**
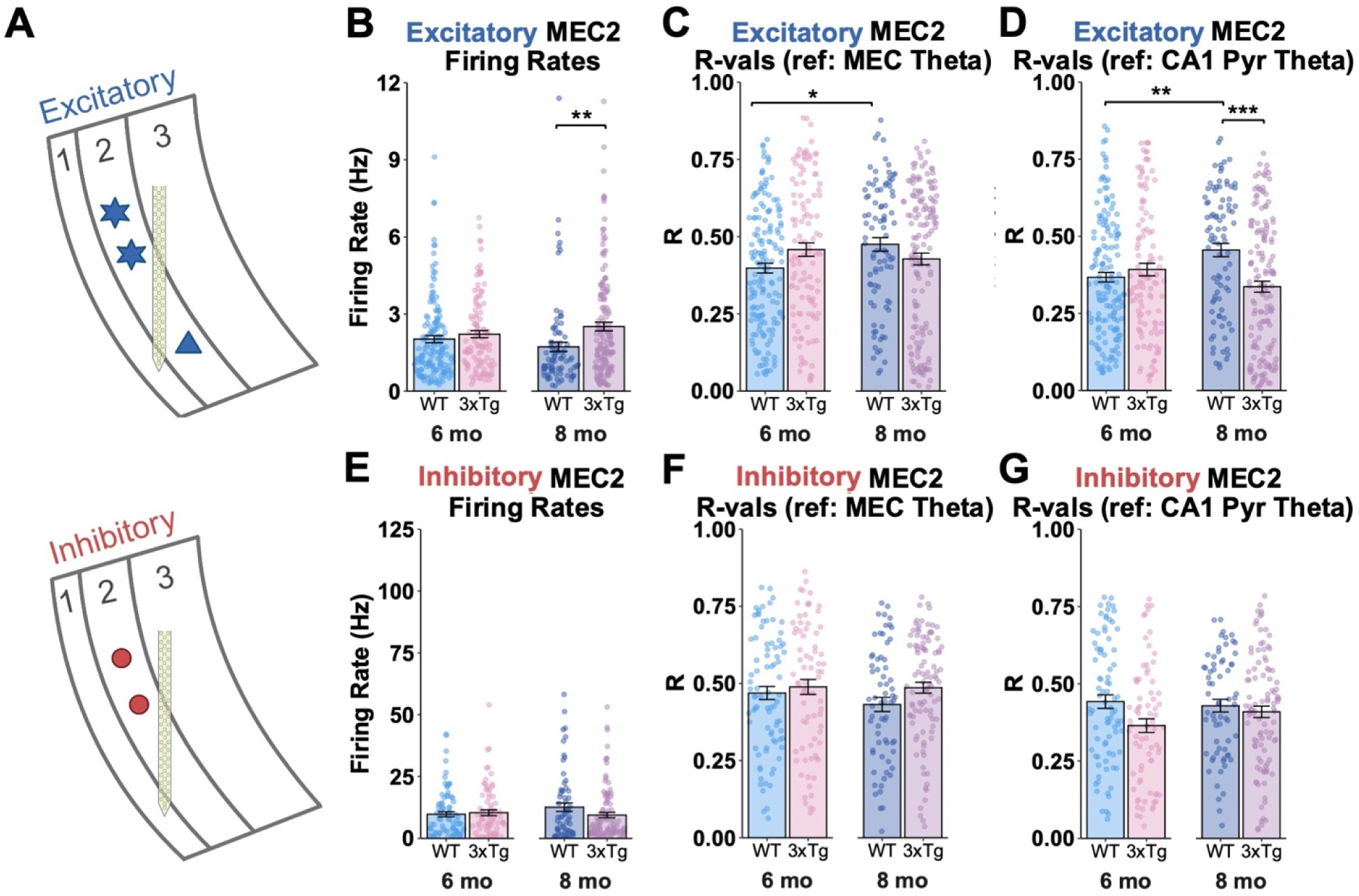
Increased excitatory firing rates and reduced phase locking to CA1 theta in MEC2 excitatory units A. Schematics showing probe location and cell types in MEC2. Excitatory stellate and pyramidal cells in MEC2 are shown in the top row in blue, while interneurons are shown in the bottom row in red. **B.** MEC2 excitatory firing rates. MEC2 putative excitatory cells in 8-month-old 3xTg mice showed increased firing rates compared to units in age-matched WT controls (3xTg 8 mo vs. WT 8 mo, Mann-Whitney (MW) test, Holm adjusted p <0.01). **C.** Phase locking strength of excitatory MEC2 units to theta oscillations recorded in MEC. R-values increased with age in WT mice (WT 6 mo vs. WT 8 mo, MW, adjusted p <0.01). **D.** Phase locking strength of excitatory MEC2 units to theta referenced in CA1 pyramidal layer. Theta phase locking increased with age in WT mice and was reduced in 8-month-old 3xTg mice relative to age-matched WT controls (MW tests, WT 6 mo vs. WT 8 mo adjusted p<0.01, WT 8 mo vs. 3xTg 8 mo adjusted p<0.001). **E.** MEC2 inhibitory firing rates. No change in firing rates of inhibitory MEC2 units across groups. **F.** Phase locking strength of MEC2 putative inhibitory neurons to theta oscillations recorded in MEC. No significant differences between groups. **G.** Phase locking strength of MEC2 inhibitory units to theta recorded in CA1. No significant differences between groups. **Excitatory Units:** WT 6 mo (n = 157), 3xTg 6 mo (n = 114), WT 8 mo cells (n = 88), 3xTg 8 mo (n = 150). **Inhibitory Units:** WT 6 mo (n = 83), 3xTg 6 mo (n = 76), WT 9 mo cells (n = 66), 3xTg 9 mo (n = 100). Extended information about # of cells per animal can be found in Supplemental Table 1. Error bars show s.e.m. *p<0.05, **p<0.01, ***p<0.001 Because single unit data did not meet ANOVA assumptions for normality and equal variances, Mann-Whitney tests were used for all comparisons (see Methods and Supplemental Table 1 for more details).

We next examined the strength of phase locking of excitatory MEC2 neurons to their local (MEC) theta oscillations, in order to assess within-MEC synchronization. MEC2 excitatory units in 3xTg and WT mice had similar phase locking to MEC theta at both time points, suggesting that synchrony within MEC2 is intact in 3xTg mice (Fig. 5C). Within the WT group, there was an increase in phase locking strength from 6 to 8-months of age, suggesting changes related to normal aging, but this increase was not detected in 3xTg mice (Fig. 5C). Though synchrony of MEC2 spike timing with local MEC activity was mostly preserved, when we examined the synchronization of MEC2 excitatory neurons with downstream CA1 theta oscillations, we found decreased phase locking in 8-month-old 3xTg animals compared to age-matched WT mice. Thus, excitatory 3xTg MEC2 neurons had reduced synchrony with downstream hippocampal theta compared to age-matched WT neurons only at our later time point, after the onset of memory impairments (Fig. 5D). Interestingly, we again observed an increase across age only in the WT group, suggesting that this deficit in 3xTg mice may emerge in part due to a failure to increase phase locking with age, rather than simply a reduction in phase-locking strength with age (Fig. 5C-D). We next extended this analysis to examine interneurons in MEC2. These interneurons provide inhibition to the surrounding excitatory cells and are also known to receive long-range input from the medial septum^82–84^ that modulates theta oscillations. We found no firing rate changes in MEC2 interneurons (Fig. 5E), and no differences in interneuron phase locking to MEC or CA1 theta at either time point (Fig. 5F-G). Overall, these results suggest a deficit in synchronization between MEC2 excitatory spiking and downstream CA1 theta in 3xTg mice that emerges across the development of memory impairments.

Given the altered MEC3 interneuron firing patterns and increased MEC2 excitatory firing rates we found in 3xTg mice, we next used immunohistochemistry to further examine MEC interneurons for signs of inhibitory dysfunction. Recent studies have highlighted vulnerability of fast-spiking interneurons across AD mouse models^23,71,85–87^. Parvalbumin expressing (PV+) inhibitory neurons provide inhibition to MEC excitatory neurons, as well as long range inhibition to hippocampus^82–84,88–91^.

These findings led us to predict that PV+ interneurons might be impacted in 3xTg mice. We found that PV+ cell counts in MEC2 and MEC3 were reduced by 6-months of age in 3xTg mice compared to WT (Fig. S8A-C). However, no changes were detectable in the hippocampus at 6 or 8 months of age (Fig. S8G-J). To determine whether reduced labeling of PV+ cells could be explained by overall neuronal loss, we also performed staining for NeuN to label all neurons. We found no differences in neuronal counts in 6-month-old 3xTg mice, suggesting that the early PV+ changes we saw were cell-type specific and could not be explained by widespread cell death (Fig. S8D-F). However, we found reductions in MEC2 and MEC3 NeuN counts by 8-months of age in 3xTg mice. Overall, MEC PV+ interneuron counts were decreased in 3xTg mice at 6-months of age, prior to any overall neuronal loss, while PV+ counts in hippocampus were unchanged at both timepoints. This suggests that PV+ interneurons in MEC are uniquely vulnerable early on in these animals. Loss or dysfunction in MEC interneurons may lead to a loss of inhibition within MEC and hippocampus, potentially contributing to hyperexcitability and altered spike timing.

### Progressive disruptions in hippocampal phase locking in 3xTg mice

Given the progressive changes in MEC-HPC synchronization between 6 and 8 months of age, we hypothesized that the timing of hippocampal neurons’ firing would also be disrupted in 8-month-old 3xTg mice. To investigate hippocampal spike timing, we examined single unit activity in CA1 and DG. We again identified putative excitatory and inhibitory units based on their waveforms, firing rates, and firing properties (Fig. S6, see Methods). The majority of CA1 units were located in and around the pyramidal layer, while the majority of DG units were located in the hilus. We first examined firing rates and found no changes between 3xTg and WT neurons in either hippocampal subregion or cell type (Fig. S9). However, when we examined phase locking of CA1 neurons, we found that phase locking to CA1 theta was reduced in both excitatory and inhibitory neurons of 3xTg mice relative to neurons from WT controls, with this deficit emerging between 6 and 8 months of age (Fig. 6A-C). These results suggest that spike timing in CA1 becomes disorganized with age in 3xTg mice. The timing of this dysfunction aligns with the onset of disrupted spike timing upstream in MEC and with broader deficits in theta synchrony and spatial memory.

**Figure 6:**
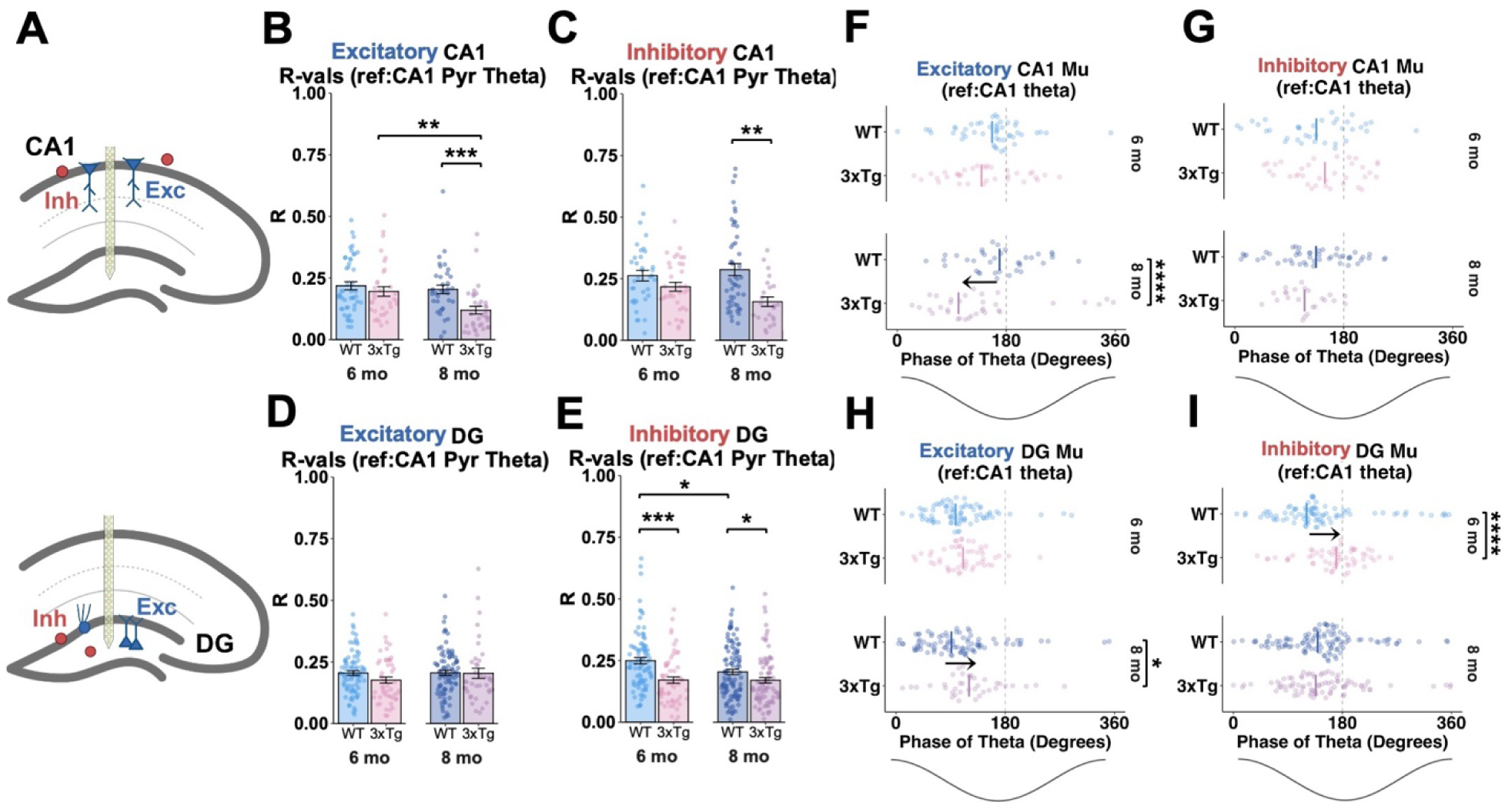
Early DG interneuron dysfunction and progressive disruptions in hippocampal spike timing in 3xTg mice. **A.** Schematics showing probe location and cell types in the hippocampus. The top row shows CA1 pyramidal cells (exc, in blue) and interneurons (inh, in red). The bottom row depicts DG hilar mossy cells and granule cells (exc, in blue - a majority of the excitatory neurons recorded here are likely hilar mossy cells), and interneurons (inh, in red). **B.** Phase locking strength of putative excitatory CA1 neurons to CA1 theta oscillations. R-values were reduced in 3xTg mice between 6 and 8 months and decreased in 8-month-old 3xTg mice compared to age-matched WTs (3xTg 6 mo vs. 3xTg 8 mo Mann-Whitney (MW) test, adjusted p = 0.003, WT 8 mo vs. 3xTg 8 mo MW adjusted p<0.001). **C.** Phase locking strength of putative CA1 inhibitory neurons to theta recorded in CA1 pyramidal layer. Phase locking of inhibitory CA1 units to local hippocampal theta was decreased in 8-month-old 3xTg mice compared to age-matched WT controls (WT 8 mo vs. 3xTg 8 mo MW adjusted p = 0.002). **D.** Phase locking strength of putative excitatory DG neurons to theta oscillations recorded in CA1 pyramidal layer. No group differences. **E.** Phase locking strength of putative DG interneurons to CA1 theta oscillations. Phase locking of DG interneurons was reduced in 3xTg mice compared to WT controls at both 6 and 8 months, as well as in normal aging (WT 6 mo vs. 3xTg 6 mo MW test adjusted p<0.001, WT 8 mo vs. 3xTg 8 mo MW adjusted p = 0.032, WT 6 mo vs. WT 8 mo MW adjusted p =0.021). **F.** Preferred firing phase of putative excitatory CA1 neurons relative to CA1 theta. 8-month-old 3xTg excitatory CA1 units shifted their theta phase earlier relative to age-matched WT control units (WT 8 mo vs. 3xTg 8 mo Kuiper test adjusted p = 0.004, Watson-Williams adjusted p< 0.0001). **G.** Preferred firing phase of putative CA1 interneurons relative to theta recorded in CA1 pyramidal layer. No differences between groups. **H.** Preferred firing phase of putative DG excitatory neurons relative to theta recorded in CA1 pyramidal layer. Schematic of theta phase is shown along the x-axis. DG excitatory units shifted their firing to a later phase of theta in 8-month-old 3xTg mice compared to age-matched WTs (WT 8 mo vs. 3xTg 8 mo, Kuiper test adjusted p<0.001, Watson-Williams test adjusted p = 0.011). **I.** Preferred firing phase of putative DG interneurons relative to CA1 theta oscillations. DG inhibitory units exhibited shifted firing phase in 6-month-old 3xTg WT mice compared to age-matched WTs (WT 6 mo vs. 3xTg 6 mo Kuiper test adjusted p<0.001, Watson-Williams test adjusted p<0.0001). **Excitatory DG:** WT 6 mo (n = 80 units), 3xTg 6 mo (n = 51), WT 8 mo (n = 90), 3xTg 8 mo (n = 38); **Inhibitory DG:** WT 6 mo (n = 88), 3xTg 6 mo (n = 58), WT 8 mo (n = 110), 3xTg 8 mo (n = 83); **Excitatory CA1:** WT 6 mo (n = 48), 3xTg 6 mo (n = 34), WT 8 mo (n = 36), 3xTg 8 mo (n = 38); **Inhibitory CA1:** WT 6 mo (n = 36), 3xTg 6 mo (n = 35), WT 8 mo (n = 49), 3xTg 8 mo (n = 23). Extended information about # of cells per animal can be found in Supplemental Table 1. Error bars show s.e.m. Lines in F-I indicate circular means. *p<0.05, **p<0.01, ***p<0.001, ****p<0.0001. Because single unit data did not meet ANOVA assumptions for normality and equal variances, Mann-Whitney tests were used for all comparisons in B, C, D, and E (see methods and Supplemental Table 1 for more details).

We also examined firing patterns in DG, which receives projections from MEC2 and sends inputs to CA1 via CA3. Here, our recorded population of excitatory DG neurons likely consists primarily of mossy cells due to their location in the hilus and relatively high firing rates^92^. Dentate granule cells, which are located in the granule cell layer and have sparse activity and low firing rates, are difficult to detect with silicon probes in awake, head-fixed animals and were rarely seen in our recordings. Though granule cells are the main recipients of input from EC2, all DG excitatory neurons reflect the timing of ongoing processing within the DG and are useful to examine the relationship between oscillatory activity and local firing in DG. We did not see significant changes in phase locking of DG excitatory neurons (Fig. 6D). Conversely, phase locking was reduced in DG interneurons recorded from 6-month-old 3xTg mice, and this decrease persisted at 8 months of age relative to neurons from age-matched WT mice (Fig. 6E). Interestingly, phase locking was also reduced in WT DG interneurons between 6 and 8 months of age, suggesting that DG interneurons are also impacted by normal early aging processes between these time points. Taken together, these findings point to both early and late-onset synchronization deficits within the hippocampus of 3xTg mice, with DG interneuron dysfunction emerging early in disease progression. Conversely, CA1 phase locking deficits emerged with a similar timeframe to the onset of memory impairment, abnormal network activity, and upstream deficits in MEC circuits.

In addition to reductions in the strength of phase locking, alterations in single unit firing patterns can also manifest as a shift in the preferred firing phase (mean firing phase, or mu-value). In WT mice, DG excitatory units tend to fire, on average, at the descending phase of CA1 theta oscillations, while CA1 excitatory units tend to fire slightly later, close to the trough (Fig. 6F,H)^68,76,78,79,93^. This relationship, where firing in DG is followed by firing in CA1 after a short delay, may facilitate and reflect information flow through the hippocampal tri-synaptic loop. Thus, a shift in spike timing in these two regions is likely to impact intrahippocampal communication. Indeed, we found a disrupted phase relationship in the timing of excitatory neurons in DG and CA1 that emerged in 8-month-old 3xTg mice. Excitatory CA1 units in 8-month-old 3xTg mice tended to fire at an earlier phase of theta relative to age-matched WT controls (Fig. 6F), while excitatory units in DG tended to fire at a later phase of theta compared to WT controls (Fig. 6H). Together, these changes both contribute to an abnormal temporal relationship between DG and CA1 excitatory neural activity and suggest that information flow through the hippocampus is disrupted in 8-month-old 3xTg mice.

In hippocampal interneurons, we found that mu values in CA1 were unaffected in 3xTg mice at either time point (Fig. 6G), while DG interneurons were shifted toward later phases of theta at 6-months in 3xTg mice (Fig. 6I). This early-onset shift in DG interneurons further suggests that 3xTg mice have early DG interneuron dysfunction, although this change was normalized by 8-months of age, perhaps due to some compensatory response to early dysfunction. Notably, none of the changes we observed in phase locking strength or preferred phase can be explained by changes in hippocampal firing rates as there were no differences in either CA1 or DG excitatory or inhibitory neurons in 3xTg mice at either time point (Fig. S9). Altogether, our analysis of phase locking in hippocampus suggests that DG interneurons may be vulnerable to early dysfunction prior to the onset of memory impairments, while CA1 dysfunction occurs later and aligns more closely with the onset of cognitive deficits.

## Discussion

By recording entorhinal-hippocampal activity at time points before or after the emergence of spatial memory impairments in 3xTg mice, we found deficits in local and long-range synchronization that coincided with the onset of spatial memory deficits at 8 months of age (Fig. 1B). We observed several network-level changes in local field potentials including reduced hippocampal theta power (Fig. 3D, F) and reduced theta coherence between MEC and CA1 in 8-month-old 3xTg mice (Fig. 3J). We also found desynchronization at the level of single units, including reduced theta phase locking of MEC3 neurons (Fig. 4), hyperactivity and impaired timing of MEC2 excitatory neuron firing relative to hippocampal theta (Fig. 5), and disrupted phase locking of downstream CA1 and DG neurons (Fig. 6). Together, these deficits in synchronization within and between MEC and hippocampus represent a disrupted circuit that is lacking the precise timing and organization that supports information processing and spatial memory in healthy animals^68,94,95^. These deficits all emerged along a similar time course to memory impairments, suggesting that they are likely contributors to the development of memory deficits in 3xTg mice.

Disrupted entorhinal-hippocampal circuit processing aligns with the time course of memory impairment in 3xTg mice

By recording neural activity at time points immediately prior to or after the onset of memory impairments, we were able to find circuit deficits that paralleled memory impairment in 3xTg mice. Many studies have examined circuit properties in mouse models of AD pathology, but most have relied on time points after the onset of memory deficits, making it difficult to determine which deficits might underlie memory impairments. Importantly, the circuit deficits that directly drive memory impairment should emerge with a similar time course to memory deficits. Conversely, early physiological changes that emerge prior to memory impairment may represent early biomarkers of disease, but are not sufficient to drive memory impairment on their own. Here, we identified several important circuit mechanisms that emerged along the time course of memory impairment, including reduced hippocampal theta power, reduced MEC-CA1 theta coherence, and disrupted spike timing of MEC and HPC neurons.

While the causal role of these synchronization processes still needs further investigation, these deficits are likely to contribute to the memory impairments observed in these animals. On the other hand, we also observed early deficits in MEC PV+ interneuron number (Fig. S8) and inhibitory spike timing in hippocampus (Fig. 6E) that emerged prior to memory impairments. These inhibitory deficits highlight interneuron vulnerability early in the disease process. While these changes are not sufficient to drive memory deficits alone, they could contribute to a cascade of changes that promote cognitive deficits later in disease progression.

Notably, the time course of memory impairment did not align with the emergence of aggregated amyloid plaques or neurofibrillary tau. While memory impairments and entorhinal-hippocampal desynchronization emerged between 6 and 8 months of age, we found little to no aggregated pathology at these time points (Fig 1C-D). Indeed, recent work extensively characterizing pathology in 3xTg mice showed an increase in insoluble Aβ in hippocampus and an onset of plaque formation in subiculum and CA1 around 12 months of age, which is in line with our immunohistochemistry results^58^ (Fig. 1D). This data suggests that non-aggregated forms of pathology are likely to drive the circuit and memory deficits we observed. Buildup of soluble APP and Aβ has been shown prior to overt plaque formation in these animals, and soluble oligomeric forms of Aβ have been shown to be particularly toxic to neurons^55,57,60,61,96–99^. Intracellular Aβ has also been found to accumulate in subiculum/CA1 prior to plaque deposition in 3xTg mice^55^. While there are no neurofibrillary tangles in 3xTg mice at the ages examined here, minimal hyperphosphorylated tau has been reported in subiculum by 6-months old, spreading into CA1 with age^55,59^. Therefore, the presence of increased APP, Aβ, and phosphorylated tau that has not yet formed plaques or tangles could contribute to the physiological changes and memory impairment we report here.

Future studies will need to examine *in vivo* physiology across mouse models with distinct mutations (i.e., only amyloid or tau related mutations) to establish which circuit changes converge or diverge across models. It is important to note, however, that the precise relationship between accumulation of pathological proteins and cognitive impairment is unclear, while changes in network activity may be more predictive of cognitive decline and more responsive to noninvasive therapies^38,100,101^. Thus, it is critical to elucidate what changes in brain activity and circuit function are most closely associated with memory impairments to design interventions that can ameliorate cognitive decline.

### Reductions in theta power and synchrony suggest impaired communication between MEC and hippocampus in 3xTg mice

We observed major deficits in the synchronization of hippocampus and medial entorhinal cortex in 3xTg mice. Because entorhinal cortex is a major driver of hippocampal theta^68,94^, reduced hippocampal theta power and CSD magnitude in 8-month-old 3xTg mice (Fig. 3) are consistent with a reduction of properly timed input from entorhinal cortex to hippocampus. This could be driven by synaptic loss or weakened synaptic transmission between entorhinal cortex and hippocampus^102–104^, and/or by disruptions in coordination between regions, an interpretation supported by the intact MEC theta power in these mice. Indeed, poor coordination of spiking between entorhinal cortex and hippocampus may impair plasticity between these regions, which could also contribute to synaptic loss and reduced hippocampal theta power.

While reduced theta power was present throughout the hippocampus (Fig. 3A), synchrony was reduced specifically between MEC and CA1, with no changes in theta synchrony between CA1 and DG (Fig. 3G-J). Prior studies have suggested a loss of synchrony between EC and hippocampus in other rodent models of AD pathology, but these findings were based on tetrode recordings from EC and CA1 pyramidal layer only, leaving uncertainty regarding when and where this desynchrony emerges and how it might relate to other neural activity patterns^14,39^. Here, the use of silicon probes allowed us to perform laminar recordings across MEC, DG, and CA1 at multiple time points, revealing a reduction of theta coherence specifically between MEC and CA1 oriens, pyramidal layer, and radiatum in 8-month-old 3xTg mice (Fig. 3J). Reduced theta synchrony between MEC and CA1 could reflect loss of synchronized inputs to these regions, such as those from medial septum, which projects to both MEC and hippocampus and modulates theta frequency^105–107^. Changes in local hippocampal interneurons or inputs from CA3 to CA1 could also impact CA1 theta oscillations, but are unlikely to fully explain the magnitude of these changes in power and coherence^68,93,108–111^. In addition, recent work has identified additional projections of MEC2 pyramidal cells to CA1^104,112^ and long range inhibitory projections from MEC to hippocampus^88,89,113^ that could also mediate MEC-hippocampal synchronization. Further work is needed to explore how these other projections within this circuitry may contribute to desynchrony in AD.

### Progressive disruptions of MEC spike timing in 3xTg mice

We also found significant disruptions in the timing of single unit activity in MEC and hippocampus. Past studies of entorhinal physiology in rodent models of AD pathology have primarily examined MEC or entorhinal cortex as a whole, without layer specificity. Here, the enhanced spatial resolution granted by silicon probes allowed us to differentiate between populations of cells in MEC2 and MEC3, which have unique projections to hippocampus. We examined theta phase locking of MEC and hippocampal units as a metric for synchrony at the single cell level. Precise spike timing is necessary for plasticity and the encoding of spatial representations, while improper timing may contribute to impaired plasticity and disrupted information coding^25–27^. In particular, precise timing of entorhinal inputs into hippocampus supports coordinated sequential firing patterns in hippocampus that have been linked to successful spatial memory encoding^7,26,75,76^.

In MEC3 neurons, we found reduced phase locking to both local and hippocampal theta in 8-month-old 3xTg mice, coincident with the emergence of memory impairments (Fig. 4). Our data indicate that MEC3 neurons are particularly vulnerable to progressive dysfunction in 3xTg mice. The loss of phase locking to both MEC and CA1 theta suggests that these changes are not driven solely by a loss of coherence between MEC and CA1, but rather to a change in the firing patterns of MEC neurons. Meanwhile, excitatory MEC2 neurons in 8-month-old 3xTg mice exhibited reduced phase locking only to CA1 theta compared to neurons from age-matched WT mice, consistent with a loss of coherence between MEC and CA1 (Fig. 5). Overall, the timing of MEC spiking relative to hippocampal theta was disrupted, suggesting that inputs to hippocampus are not arriving at the optimal time, which could drive impaired plasticity, spatial processing, and spatial memory. Impaired timing of MEC units may also contribute to the progressive changes in phase locking we found in CA1, where projections from MEC terminate.

### Hyperexcitability and interneuron vulnerability in MEC of 3xTg mice

In addition to poor timing, inputs from MEC to hippocampus could also be disrupted by a loss of excitatory-inhibitory (E/I) balance in MEC. MEC hyperexcitability in 3xTg mice has been previously characterized *in vitro* by our lab and others^22,80^. We recently showed increased firing and reduced E/I balance specific to MEC2 stellate cells (but not other excitatory MEC2/3 cell types) in 10-month-old 3xTg mice *in vitro* ^22^. Here, we found an increase in firing rates of 3xTg excitatory MEC2 units at 8 months relative to age-matched WT units (Fig. 5B). Though we are unable to differentiate between stellate and pyramidal neurons in MEC2 *in vivo*, our past work *in vitro* suggests that this is driven primarily by stellate cells, which project to the dentate gyrus. Broadly, Aβ has been shown to promote increased excitability and/or hyperactivity in neurons by various mechanisms including disrupting glutamate reuptake, even without plaque formation^96,97,99,114^. Thus, it is possible that these changes are due to the presence of increased Aβ. However, other work has suggested that impaired inhibition may also contribute to or exacerbate increased firing in excitatory cells^22,23,115^. Notably, we did not see changes in firing rate in any hippocampal cell types (Fig. S9), suggesting a specific vulnerability to hyperactivity in MEC.

Local PV+ interneurons provide a major source of inhibition to excitatory cells in MEC and hippocampus^83,84,93,116,117^. Hippocampal interneuron dysfunction has been widely characterized in rodent models of AD pathology^8,71,85,87,118–121^, while entorhinal interneuron vulnerability has gained interest more recently^23^. Here, we found reduced PV+ interneuron labeling in MEC of 3xTg mice by 6-months of age (Fig. S8A-C). This likely reflects cell death of PV+ interneurons, but could also be due to dysfunction of these cells that leads to reduced parvalbumin expression. Since we found no changes in NeuN labeling in 6-month-old 3xTg mice (Fig. S8D-F), PV+ neurons appear to have a unique early vulnerability that precedes any widespread neuronal death. Overall, impaired inhibition and increased excitatory firing rates could promote excitotoxicity, potentially leading to neuronal loss in MEC and loss of inputs from MEC to hippocampus by 8 months of age. PV+ inhibitory neurons in MEC also receive input from medial septum that is critical for theta generation, and send long range inputs to CA1, making them a point of interest for future work investigating the mechanisms underlying synchrony in these circuits.

### Progressive CA1 phase locking deficits and early DG interneuron dysfunction in 3xTg mice

CA1 serves as an integration site for information received directly from EC3 and indirectly from EC2 via DG/CA3. We found reductions in phase locking strength of CA1 excitatory and inhibitory units in 3xTg mice that primarily emerged at 8-months of age (Fig. 6B-C). These changes could be linked to the upstream changes we see in MEC firing patterns or to dysfunction of local interneurons, which are known to gate entorhinal inputs and modulate spiking of excitatory cells^93,122,123^. Altered firing patterns in CA1 could disrupt the formation and maintenance of spatial maps, thereby contributing to spatial memory impairments. Indeed, previous work has shown worse spatial coding in CA1 place cells in 3xTg mice^13^. We also examined phase locking earlier in the hippocampal loop in DG, which receives input from MEC2. We have previously shown that theta phase locking of DG interneurons is disrupted in a mouse model of chronic temporal lobe epilepsy with significant spatial memory deficits and disrupted hippocampal spatial coding^124,125^. Here, we find early deficits in DG interneuron phase locking strength in a mouse model of AD pathology (Fig. 6E), as well as a reduction in phase locking across normal aging. Thus, changes in interneuron phase locking may be a common mechanism of hippocampal circuit dysfunction across neurological disorders and aging.

Lastly, we examined the preferred firing phase of hippocampal units. In WT mice, DG excitatory units tended to fire at the descending phase of theta oscillations, while CA1 units fired slightly later, at the trough^68,76,78,79,93^. This time lag likely reflects synaptic transmission through the tri-synaptic loop, from DG to CA1 via CA3. In 3xTg mice, however, CA1 units shifted their firing earlier, while DG units shifted their firing later, thereby bringing the average firing phase of these populations closer together (Fig. 6F, H). Previous work has shown that the theta phase preference of CA1 and DG excitatory units depends upon the relative strength and timing of entorhinal and intrahippocampal inputs^7,76^. Based on this evidence, the observed shifts in spike timing may reflect weakening or impaired timing of MEC inputs to CA1 and DG. This could result in an increased influence of other inputs from CA3 to CA1 and from commissural pathways to DG. These shifts toward earlier CA1 and later DG spike timing could also reflect subtle hyper- and hypo-excitability in these two regions, respectively. However, since we do not see any changes in firing rates, a more likely explanation is that the strength and/or timing of the inputs that these cells receive are altered.

### Direct manipulations of entorhinal-hippocampal synchrony are needed to elucidate causal relationships between synchrony and memory

Here, we identified several circuit deficits that align with the onset of memory impairments in 3xTg mice. These results generate a new hypothesis that the disrupted timing of entorhinal-hippocampal neural activity directly drives memory impairment. However, while the onset of spatial memory impairments aligns with the emergence of impaired theta power and synchrony, this finding is still correlational and thus not sufficient to establish a causal link. Fortunately, new technologies are being developed to enable precise manipulations of neural synchrony between brain regions as well as precise manipulations of the phase locking of neurons to endogenous theta oscillations^126,127^. Using such tools, it will be possible to test hypotheses about the causal role of synchrony in spatial memory and examine the relationships between abnormal spike timing and theta synchrony. Future work will need to directly test whether manipulations such as restoring theta synchrony between MEC and CA1 or restoring the spike timing of MEC3 neurons relative to hippocampal theta could improve circuit function and rescue impairments in spatial coding and spatial memory.

## Methods

### Key Resources Table

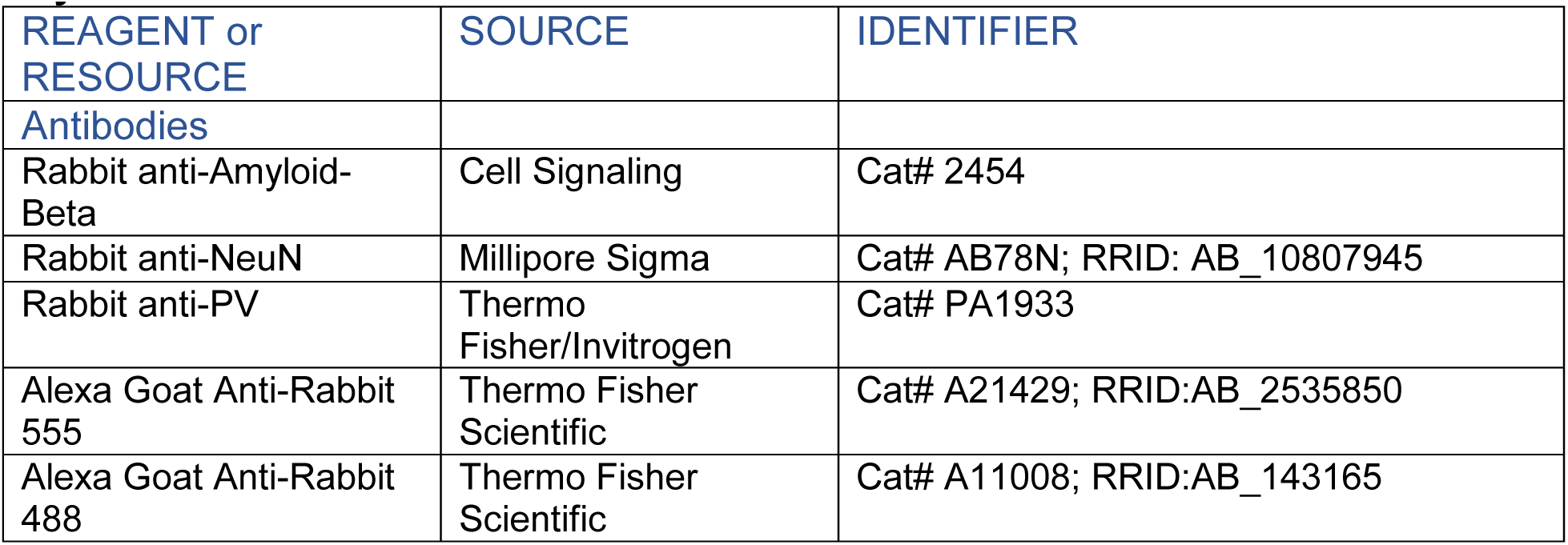
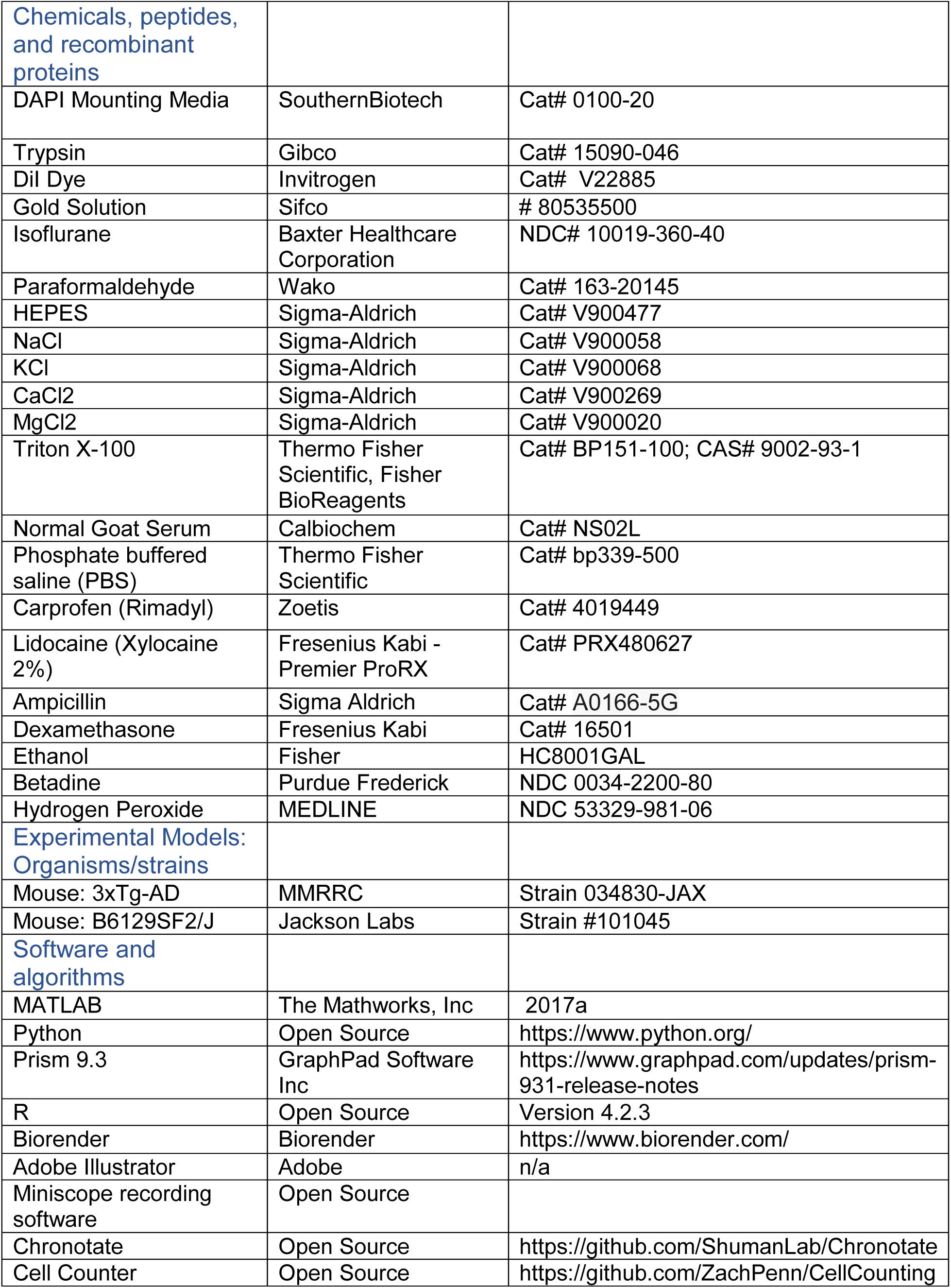

| REAGENT or RESOURCE | SOURCE | IDENTIFIER |
| --- | --- | --- |
| **Antibodies** |  |  |
| Rabbit anti-Amyloid-Beta | Cell Signaling | Cat# 2454 |
| Rabbit anti-NeuN | Millipore Sigma | Cat# AB78N; RRID: AB_10807945 |
| Rabbit anti-PV | Thermo Fisher/Invitrogen | Cat# PA1933 |
| Alexa Goat Anti-Rabbit 555 | Thermo Fisher Scientific | Cat# A21429; RRID:AB_2535850 |
| Alexa Goat Anti-Rabbit 488 | Thermo Fisher Scientific | Cat# A11008; RRID:AB_143165 |
| Chemicals, peptides, and recombinant proteins |  |  |
| DAPI Mounting Media | SouthernBiotech | Cat# 0100-20 |
| Trypsin | Gibco | Cat# 15090-046 |
| Dil Dye | Invitrogen | Cat# V22885 |
| Gold Solution | Sifco | # 80535500 |
| Isoflurane | Baxter Healthcare Corporation | NDC# 10019-360-40 |
| Paraformaldehyde | Wako | Cat# 163-20145 |
| HEPES | Sigma-Aldrich | Cat# V900477 |
| NaCl | Sigma-Aldrich | Cat# V900058 |
| KCl | Sigma-Aldrich | Cat# V900068 |
| CaCl<sub>2</sub> | Sigma-Aldrich | Cat# V900269 |
| MgCl<sub>2</sub> | Sigma-Aldrich | Cat# V900020 |
| Triton X-100 | Thermo Fisher Scientific, Fisher BioReagents | Cat# BP151-100; CAS# 9002-93-1 |
| Normal Goat Serum | Calbiochem | Cat# NS02L |
| Phosphate buffered saline (PBS) | Thermo Fisher Scientific | Cat# bp339-500 |
| Carprofen (Rimadyl) | Zoetis | Cat# 4019449 |
| Lidocaine (Xylocaine 2%) | Fresenius Kabi - Premier ProRX | Cat# PRX480627 |
| Ampicillin | Sigma Aldrich | Cat# A0166-5G |
| Dexamethasone | Fresenius Kabi | Cat# 16501 |
| Ethanol | Fisher | HC8001GAL |
| Betadine | Purdue Frederick | NDC 0034-2200-80 |
| Hydrogen Peroxide | MEDLINE | NDC 53329-981-06 |
| Experimental Models: Organisms/strains |  |  |
| Mouse: 3xTg-AD | MMRRC | Strain 034830-JAX |
| Mouse: B6129SF2/J | Jackson Labs | Strain #101045 |
| Software and algorithms |  |  |
| MATLAB | The Mathworks, Inc | 2017a |
| Python | Open Source | https://www.python.org/ |
| Prism 9.3 | GraphPad Software Inc | https://www.graphpad.com/updates/prism-931-release-notes |
| R | Open Source | Version 4.2.3 |
| Biorender | Biorender | https://www.biorender.com/ |
| Adobe Illustrator | Adobe | n/a |
| Miniscope recording software | Open Source |  |
| Chronotate | Open Source | https://github.com/ShumanLab/Chronotate |
| Cell Counter | Open Source | https://github.com/ZachPenn/CellCounting |

### Experimental Model and Subjects

All experiments were performed in accordance with Icahn School of Medicine at Mount Sinai’s Institutional Animal Care and Use Committee (IACUC) and the US National Institutes of Health (NIH) guidelines and protocols. Male and female 3xTg (MMRRC 034830-JAX) ^51^ and wild-type (WT; B6129SF2/J, strain #101045, The Jackson Laboratory) were used in all experiments. Breeders were purchased from the Mutant Mouse Resource Center (MMRRC) and Jackson Laboratories, and all experimental animals were bred in breeding facilities at Mount Sinai. Mice received *ad libitum* food and water unless water restricted for virtual reality training. All animals were kept on a 12-hour light/dark cycle (lights on at 7:00AM) and housed with at least one cage-mate. If a socially compatible littermate was not available, animals were housed with an ovariectomized female mouse (129X1/SvJ, Jackson Laboratory). Experiments took place during the light phase.

### Novel Object Location Behavior

Novel object location behavior was performed in 6 and 8-month-old WT and 3xTg mice. Average age for each group: 6.07 +/- 0.03 (WT 6 mo), 5.87 +/- 0.10 (3xTg 6 mo), 8.05 +-/ 0.11 (WT 8 mo), 8.10 +/- 0.14 (3xTg 8 mo). Animals were handled 1x daily in the experimental room for 5 days, followed by two 5-minute-long sessions of habituation to the behavioral chambers across two days with no objects present. Each chamber consisted of a white 1 ft x 1ft x 1ft white plastic box with spatial cues on each interior wall. On the following day, mice were placed in the chamber with two identical objects (plastic toys taped to the floor at two corners of the box) and allowed to explore until they reached a criterion of 30s of total exploration time. This criterion was used to ensure that all groups explored the objects equally and thus had the opportunity to learn about their locations, despite potential differences in activity levels between groups^128^.

Exploration was defined as time the mouse spent within 2 cm of the object, with its nose facing the object, and actively sniffing the object. After reaching this criterion, mice were removed from the chamber and returned to their home cage. Four hours later, animals were placed back in the same chamber with one object moved to a new location and allowed to explore for 20 minutes. Animals that did not explore for at least 30s within 20 minutes during training, that met criterion during the training phase in under 5 minutes, or that had a bias during training such that they spent 70% or more of their exploration time with one of the two objects were excluded from the experiment. Videos were scored using Chronotate^129^, by an experimenter blinded to experimental group. Only the first 20s of exploration time during the test session was scored. Performance was calculated by taking time spent exploring the novel object divided by the total time exploring both objects.

### Immunohistochemistry

Mice were perfused with phosphate buffered saline (PBS, Fisher BP399), followed by 4% paraformaldehyde (Wako). Brains were post-fixed in 4% paraformaldehyde overnight, followed by 30% sucrose (Fisher). After saturation (∼ 2 days), they were frozen at −80 degrees Celsius until sectioning. Brains were sectioned sagittally on a cryostat at 40 µm thickness, and sections were stored in 1x PBS for up to 1 day prior to staining. Sections were washed 3x for 10 minutes in TBS. This was followed by two hours of blocking at room temperature with 0.3% Triton-X-100 and 3% Normal Goat Serum in TBS. Next, sections were incubated in 3% normal goat serum with 0.3% Triton-X-100 in TBS and primary antibody (NeuN, anti-rabbit Millipore Sigma ABN78, 1:2000, or PV anti-rabbit, ThermoFisher PA-1933, 1:1000) overnight on a shaker at 4 degrees Celsius. On the following day, sections were washed 4x for 15 minutes in 1x TBS, before being incubated with secondary antibody for 2 hours at room temperature (Goat anti-rabbit Alexa Fluor 488, 1:500). Finally, sections were washed again 4x for 15-minutes in PBS.

For Aβ immunohistochemistry, an additional step was added at the start of the protocol to improve antigen retrieval^130^. Plates were incubated on a heating block at 80 degrees Celsius for 45 minutes in sodium citrate buffer. Tissue was then incubated with the primary antibody (Aβ Cell Signaling #2454, 1:400) for 48 hours.

Following staining, tissue was mounted on microscope slides with DAPI mounting media (SouthernBiotech DAPI Fluoromount-Gm, 0100-20). Slices were imaged with a 20x objective using a Leica DM6B fluorescence microscope with a Lumencor Light Engine and Leica DFC9000 GT camera. PV and NeuN Cell counting were performed using a cell counting algorithm in Python (https://github.com/ZachPenn/CellCounting) by an experimenter blinded to experimental group. For all quantitative experiments, imaging parameters were maintained across subjects. Regions of interest (ROIs) were drawn manually around each layer or subregion by an experimenter blinded to group and counts for a given slice and region were divided by the total area of the corresponding ROI to generate counts per square mm.

### Headbar Surgery

All animals used in electrophysiology experiments underwent stereotaxic surgery to attach a stainless steel headbar to the skull. Mice were anesthetized with 1-3% isoflurane at ∼4.5 or ∼6.5 months of age and fixed in a stereotaxic surgical apparatus. Lidocaine (2%, ∼0.05ml, Fresenius Kabi) was administered subcutanerously around the epaxial muscles. Hair was removed and skin was cleaned with 70% ethanol (Fisher) and betadine (Purdue Frederick). The skull was exposed and the top layer of epaxial muscles were detached from the skull using fine forceps. Hydrogen peroxide (3%, MEDLINE) was used to remove the fascia and allow clearer identification of bregma and lambda. The exposed skull was scored with a scalpel. The skull was aligned stereotaxically prior to attaching the headbar to the skull using cyanoacrylate glue and dental cement (LANG Dental). A barrier around the exposed skull was created by building up a well of dental cement. The well was filled with Kwik-Sil (World Precision Instruments, Cat# 600022) and covered with a final thin layer of dental cement. Subcutaneous carprofen (5 mg/kg, Zoetis) was delivered subcutaneously during and for two days following surgery. Ampicillin (20 mg/kg, Sigma Aldritch) was delivered during and for six days following surgery. Animals recovered on a heating pad before being returned to their housing facility.

### Virtual Reality Training

Animals were handled for 4 minutes per day for at least 3 days, then habituated to head-fixation on a flat surface for at least 3 days or until they were able to walk calmly while head-fixed. Beginning during the handling phase, mice were water-restricted and weighed daily to ensure that they maintained ∼82% of their starting body weight. Additional water was provided if animals showed signs of dehydration or weight dropped below 78%. Next, animals were head-fixed and secured on top of a Styrofoam ball and allowed to habituate to the ball for ∼30 minutes per day for 3-4 days. Once animals were able to walk comfortably on the ball, they were trained to lick a water port to receive 4 μL drops of water while headfixed. Our lickport uses an infrared beam to detect licks and deliver water via a solenoid valve (Lee Company) during lick training ^131^. Once mice learned to lick readily for water (∼3-4 days), animals were introduced to a virtual reality (VR) environment designed in MATLAB using the open-source software, ViRMEn ^132^, where they were required to run to the end of a track to receive a 4uL water rewards, after which they were automatically teleported back to the start of the track. Running speed was measured using an optical motion tracker located along the underside of the Styrofoam ball. The reading from this sensor triggered corresponding movement of the VR environment. The virtual linear track was displayed on three 24-inch computer monitors angled around the Styrofoam ball and centered slightly above the mouse’s eye-line. Once mice were running 100 trials within a 1-hour session they were considered ready for recording. On average, VR training took 8-10 days.

### Craniotomy and Ground Wire Implantation Surgery

On the day before recording, a craniotomy surgery was performed to expose the brain over the desired recording sites. The dental cement and Kwik Sil covering the skull were removed and a burr hole was drilled over the left cerebellum and an Ag/AgCl coated wire (Warner Instruments) was slipped between the skull and dura to serve as a reference. The wire was secured in place with cyanoacrylate glue and dental cement. Two craniotomies were drilled over right hippocampus and entorhinal cortex, respectively, each around 1.6 mm in diameter, centered around the desired recording coordinates. Craniotomies were covered with buffered artificial cerebrospinal fluid (ACSF; in mM: 135 NaCl, 5 KCl, 5 HEPES, 2.4 CaCl2, 2.1 MgCl2, pH 7.4)^124,133^.The skull was once again covered with Kwik-Sil and animals were allowed to recover on a heating pad overnight prior to recording. Carprofen (5 mg/kg) and dexamethasone (0.2 mg/kg, Fresenius Kabi), were given subcutaneously following surgery to reduce pain and inflammation.

### Acute Silicon Probe Recordings

Silicon probes were obtained through the Masmanidis Lab at UCLA, and their production and distribution were supported by the NSF NeuroNex program (more information available at: https://masmanidislab.neurobio.ucla.edu/images/microprobe_info.pdf). Two 256-channel silicon probes were used in all experiments, with one probe inserted into hippocampus and the other inserted into MEC. Each probe consisted of 4 shanks with 64 channels per shank. Recording sites spanned across the bottom 1.050mm of the shank across three columns with 25 µm vertical spacing and 20 µm horizontal spacing between channels. Probes were electroplated with gold solution (Sifco) to achieve 0.1 to 0.5 MΩ impedance. Prior to recording, probes were coated in Di-I to facilitate histological verification of probe locations.

On the recording day, the mouse was head-fixed on the Styrofoam ball as usual. Kwik-Sil over the skull was removed and the exposed brain was covered with ACSF. The skull was aligned to sit flat to ensure accuracy of probe targeting, and the ground wire was connected to the amplifiers to serve as a reference. The entire behavioral and recording setup are secured to an air table to reduce the potential influence of any external vibrations on electrophysiological signals. Probes were aligned to bregma and inserted into the brain at roughly the following coordinates: Hippocampus: 1.8mm posterior, 1.35mm right, 2.2mm ventral from bregma or until all hippocampal layers were detected upon examination of physiological signals; MEC: 4.9mm posterior, 3.1mm right, and lowered until the bottom of the probe appeared to reach layer 1 (indicated by a shift in theta phase and reduced spiking activity). Micromanipulators (Sutter Instruments) were used to lower probes into the brain at a rate of 2-5 µm/sec, which has been shown to improve cell yield in silicon probe recordings^134^. After reaching the desired recording location, probes were left to settle in the brain for at least one hour prior to beginning recording.

Signals were sent to four Intan headstages (RHD 128-Channel Recording Headstage, Intan Technologies, two connected to each probe). An Intan recording controller (RHD2000 Intan 1024ch Recording Controller, Intan Technologies) was used to collect the signals at 25kHz, along with analog signals reflecting the position of the animal on the linear track, water delivery from the lickport, licks sensed by an infrared beam break in front of the water port, and running speed (measured via an optical sensor that recorded ball rotation speed). Recordings lasted between 1.5-4 hours. Average ages in months by group at time of recording: 6.19 +/- 0.16 (WT 6 mo), 6.31 +/- 0.16 (3xTg 6 mo), 8.36 +/- 0.18 (WT 8 mo), and 8.41 +/- 0.28 (3xTg 8 mo).

### Histology

Following recording, silicon probes were removed from the brain. Mice were perfused with 1x PBS followed by 4% paraformaldehyde. Brains were post-fixed in 4% paraformaldehyde overnight, followed by 30% sucrose. After saturation, they were frozen on dry ice and kept frozen at −80 degrees until sectioning. Brains were sectioned sagittally on a cryostat at 80 µm thickness, and sections were stored in 24 well plates in 1x PBS. Immunohistochemistry for NeuN was performed to aid in identifying cell layers in MEC. Procedures for staining were identical to those described above in the “Immunohistochemistry” section, with one exception.

Triton was replaced by Tween for this tissue to maintain presence of Di-I in the slices. Locations of recording channels were determined by visualizing the Di-I labeled probe tracts for each animal and cross-referencing with the corresponding physiological signals. Locations of theta phase shifts, peak theta power, peak fast ripple power, relative changes in spiking activity, and overall coherence patterns across channels were used to estimate where each channel on a given shank was located^75,78,124,135,136^.

MEC layer assignments were made using the theta phase shift in layer 1-2, the relative increase in spiking activity in MEC2 compared to MEC1 and 3, and in some cases the increased phase locking of neurons in MEC2 relative to MEC3^78^. For hippocampus, CA1 the mid-point of the pyramidal layer was defined as 1-2 channels before the initiation of a theta phase shift and confirmed by checking for the presence of a peak in fast ripple power. The boundary between radiatum and lacunosum moleculare was defined by the end of the theta phase shift. The boundary between lacunosum moleculare and molecular layer was defined by the presence of a peak in theta power. Remaining hippocampal layer boundaries were defined by plotting coherence across all channels and looking for blocks of high coherence that occur within a layer.

### Analysis of Local Field Potential and Single-Unit Data

All data preprocessing was performed using custom scripts in MATLAB 2017A. First, data files were concatenated to form a single file for each channel. For local field potential (LFP) analysis, data was downsampled to 1000 Hz and bandpass filtered for more specific frequency bands as needed. For LFP analyses, one shank from MEC and one shank from hippocampus were used for each animal. The shank with the best coverage of the target regions and the fewest bad channels (i.e. channels with no signal or poor impedance) was chosen for analysis. To calculate theta power and coherence, data was first filtered for oscillations in the theta frequency band (5-12 Hz). Next, the amplitude of theta oscillations over time was calculated by taking the envelope of the theta filtered signal. This amplitude was then averaged across time, only including time points during which animals were running within a range of speeds (see Fig. S2). This range was selected to ensure that there were no differences in running speed across groups within the analyzed recording periods (Fig. S2C). This subsampling approach was used for all LFP and single unit analyses to ensure that our results could not be explained by differences in running speed or time spent running across groups. Coherence of theta filtered LFPs was calculated using an adapted version of the coherency function in the Chronux library^137^ and averaged across frequencies between 5-12 Hz. Coherence was calculated across the recording in one second bins and then subsampled to include only time periods where animals were running at speeds within the pre-defined thresholds.

For power calculations, power was averaged across all channels within a layer for each animal. To generate coherence values for a given pair of layers, coherence between each potential pair of channels was calculated and the resulting matrix was averaged. For frequency calculations in Fig S5 a wavelet transformation was used to calculate power spectral density between 5-12 Hz. For phase locking analyses, a channel in mid CA1 pyramidal layer prior to any theta phase shift was used as the CA1 theta reference and a channel in mid-MEC2 was used as the MEC theta reference. These reference channels were checked manually to ensure they were free from noise or any obvious artifacts.

Current spectral density (CSD) was calculated using code obtained from the Buzsáki lab repository (https://github.com/buzsakilab/buzcode). By taking the second spatial derivative of the LFP data across channels, we were able to isolate local current sources and sinks that were unlikely to be generated by volume conduction. This allows more precise localization and quantification of currents generated within the hippocampus. One row of 21 linear channels (each silicon probe shank contained 3 rows with 64 channels total) was used for analysis, and average CSDs were calculated across all theta cycles that occurred while an animal was running within the speed thresholds.

For single unit analysis, data from each recording was background subtracted and converted to a binary file. Kilosort 2.5 was used for initial spike sorting, followed by manual curation in Phy^138–140^. An experimenter blinded to experimental group manually inspected all units to ensure that each unit had a plausible waveform (i.e. not noise) and low refractory violations. Any units with less than 20 spikes on during periods where the animal was running within the speed threshold were excluded from further analysis.

Because our recordings lasted an hour or more, a given cell may only be detectable for a fraction of this time. Thus, calculating an average firing rate across the full length of the recording may not be an accurate reflection of a cell’s firing activity. To account for this, we first divided the recording into one-minute bins and calculated each cell’s firing rate in each, as well as a mean firing rate across all bins. Any bins where the firing rate was less than 10% of the mean firing rate were excluded from further analysis. This allowed us to keep long periods where a cell was inactive or not detectable from influencing mean firing rate calculations.

Phase locking and firing rate analysis was again restricted to spikes that occurred while animals were running within the speed threshold. Because firing rates and phase locking may differ between running and not running periods, or based on running speed, this again allows us to exclude the possibility that differences between animals/groups are due to differences in running behavior. Phase locking was measured with both R-values (precision of phase preference) and mu-values (mean phase of firing) relative to theta oscillations. For each animal, we performed a Hilbert transformation on the theta filtered LFP data and found the locations of theta peaks(0°)/troughs(180°). For each unit, we found the phase of theta at each spike time and used the circular statistics toolbox in Matlab to calculate the mean resultant vector length (R) and the mean direction (mu) for each unit.

### Quantification and Statistical Analysis

All statistics were performed in R version 4.2.3 or GraphPad Prism 9.3.1. The Holm method was used to control for multiple comparisons unless otherwise specified. All test statistics, p-values and information about multiple comparison corrections can be found in Supplemental Table 1. Behavior and cell counting or plaque counting data were analyzed using two-way Genotype x Age type III ANOVA. Posthocs were performed using the emmeans package in R to calculate t statistics. Only posthoc comparisons between genotypes within a given age or between ages within a given genotype were performed.

Power and coherence data were analyzed with two-way Genotype x Layer repeated measures type III ANOVA using the ezANOVA function from the R ez package, followed by posthoc t-tests for each layer where applicable. Due to some animals missing data for some layers in the CSD analysis and in the MEC power analysis, these data sets were analyzed using mixed effects models.

Our single unit firing rate and R-value data did not meet the ANOVA assumptions of normality and equal variance across groups, so for these analyses the Mann-Whitney test was used to compare groups. Only comparisons between genotypes within a given age or between ages within a given genotype were performed.

For mu-value data, we used the circular package in R. The Kuiper test was first used to determine whether distributions of mu values differed between groups. If the Kuiper test was significant, an equal kappa test was performed to determine whether groups differed in their concentrations. If no differences in concentration were present, we then used the Watson-Williams test to ask if circular means differed between groups. Genotypes were compared at each age and the Bonferroni method was used to control for multiple comparisons.

Unless otherwise stated, all data are presented as mean +/- SEM, where n represents the number of cells and N represents the number of animals. In the case of mu values, circular means are shown. All statistical tests with post hoc correction were two-tailed with thresholds of significance set at p < 0.05. P-values less than 0.05 are denoted with asterisks, and all posthoc tests with p-values between 0.05 and 0.075 are noted explicitly in figure legends. Additional details of all statistical tests can, including all nonsignificant p-values, can be found in Supplemental Table 1.

## Supporting information

Supplemental Table 1 - Statistics

## Acknowledgements

We thank all members of the Cai and Shuman labs for their thoughtful feedback throughout this project. We also thank Drs. Ana Pereira and Joe Castellano for their insights into AD pathology and immunohistochemistry experiments. This work was supported by T32AG049688 and F31AG069496 to LMV, R03NS111493, RF1AG072497, R01NS116357, and R01NS136590 to TS, F31NS134301 to PAP, American Epilepsy Society Predoctoral Fellowship to YF, F32NS116416, AES Postdoctoral Fellowship, and Simons Collaboration on Plasticity and the Aging Brain Transition to Independence Award to ZCW, and R01MH120162, DP2MH122399, and R56MH132959 to DJC. Thank you to Sotiris Masmanidis and the Masmanidis lab for supplying our silicon probes with the support of the NSF NeuroNex program, Award #1707408. Finally, we thank BioRender for providing resources to streamline figure creation and Jill Gregory for assistance with the schematic in Fig. 2B.

---

**Figure S1 [Related to Figure 2].**
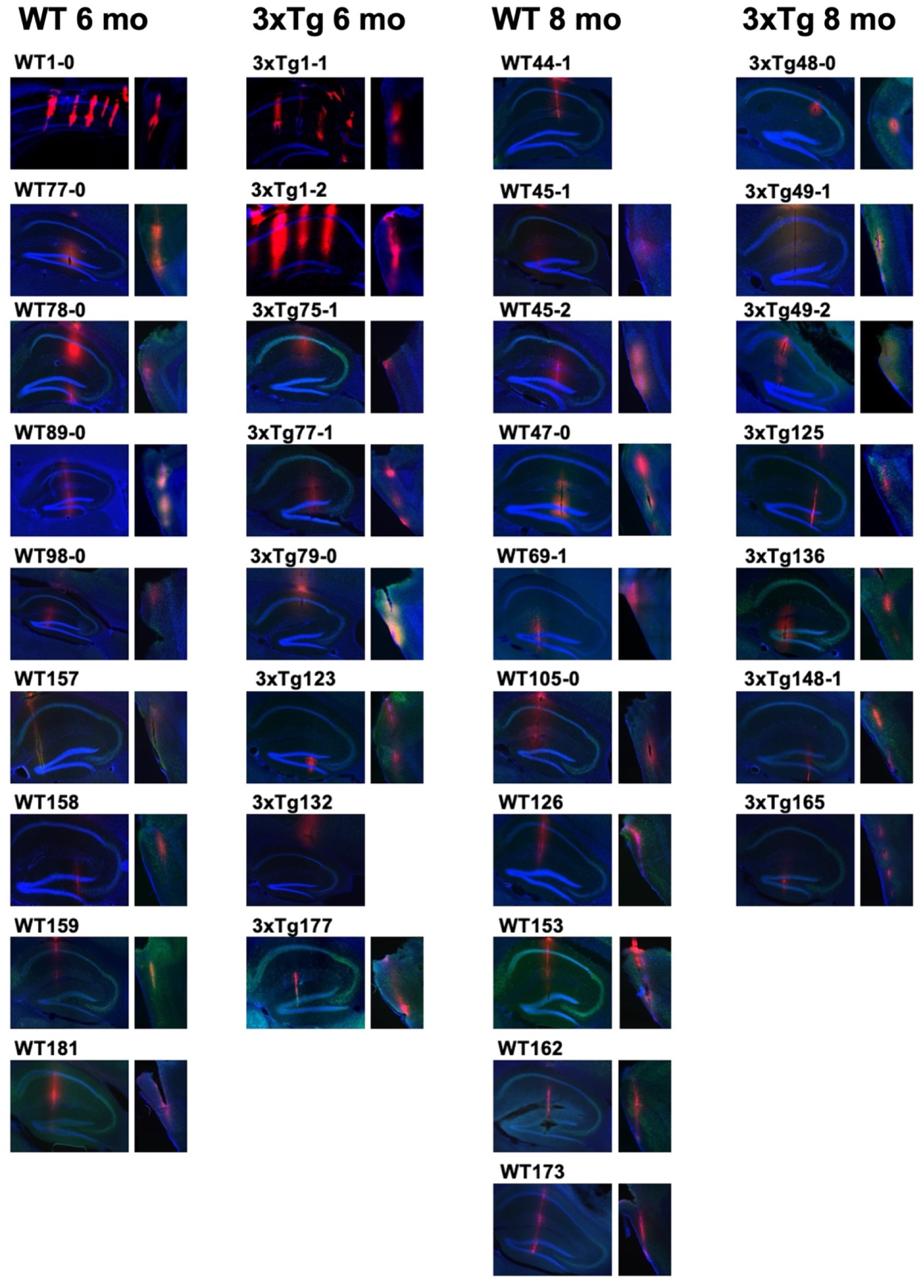
Silicon probe histology. Probes were painted with Di-I prior to recording to facilitate later verification of probe locations. Brains were sliced sagittally such that a single shank is visible in a given slice (exceptions: hippocampi from WT1-0, 3xTg1-1, and 3xTg1-2 were sliced coronally). One shank in MEC and one shank in hippocampus are shown for each animal. These shanks correspond to those used for LFP analysis in each animal. Missing MEC images indicate that data for MEC was not available for that animal.

**Figure S2 [Related to Figure 3].**
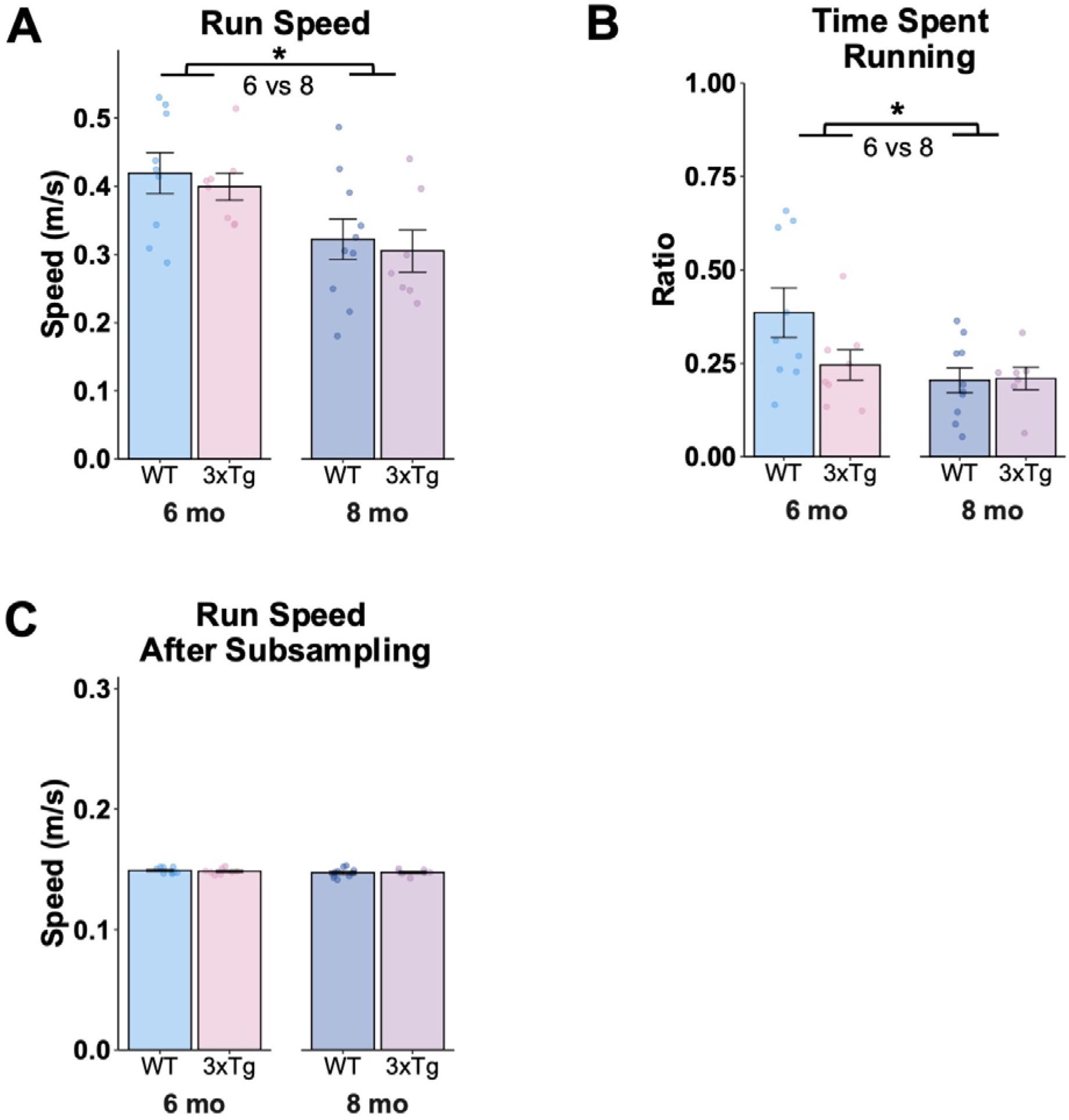
Subsampling to account for differences in running behavior across groups. **A.** Running speed by group, averaged across all times when animals were running. 8-month-old animals had a slower running speed on average (Two-way ANOVA, main effect of age, F(1, 30) = 11.04, p=0.002). **B.** Percent of recording time spent running. Aged animals spent a smaller proportion of the recording time running (Two-way ANOVA, main effect of age, F(1, 30) = 5.42, p = 0.027). **C.** Running speed after subsampling for periods where animals were running within a narrow range of speeds. No differences across groups. Error bars show s.e.m. *p<0.05, **p<0.01.

**Figure S3 [Related to Figure 3].**
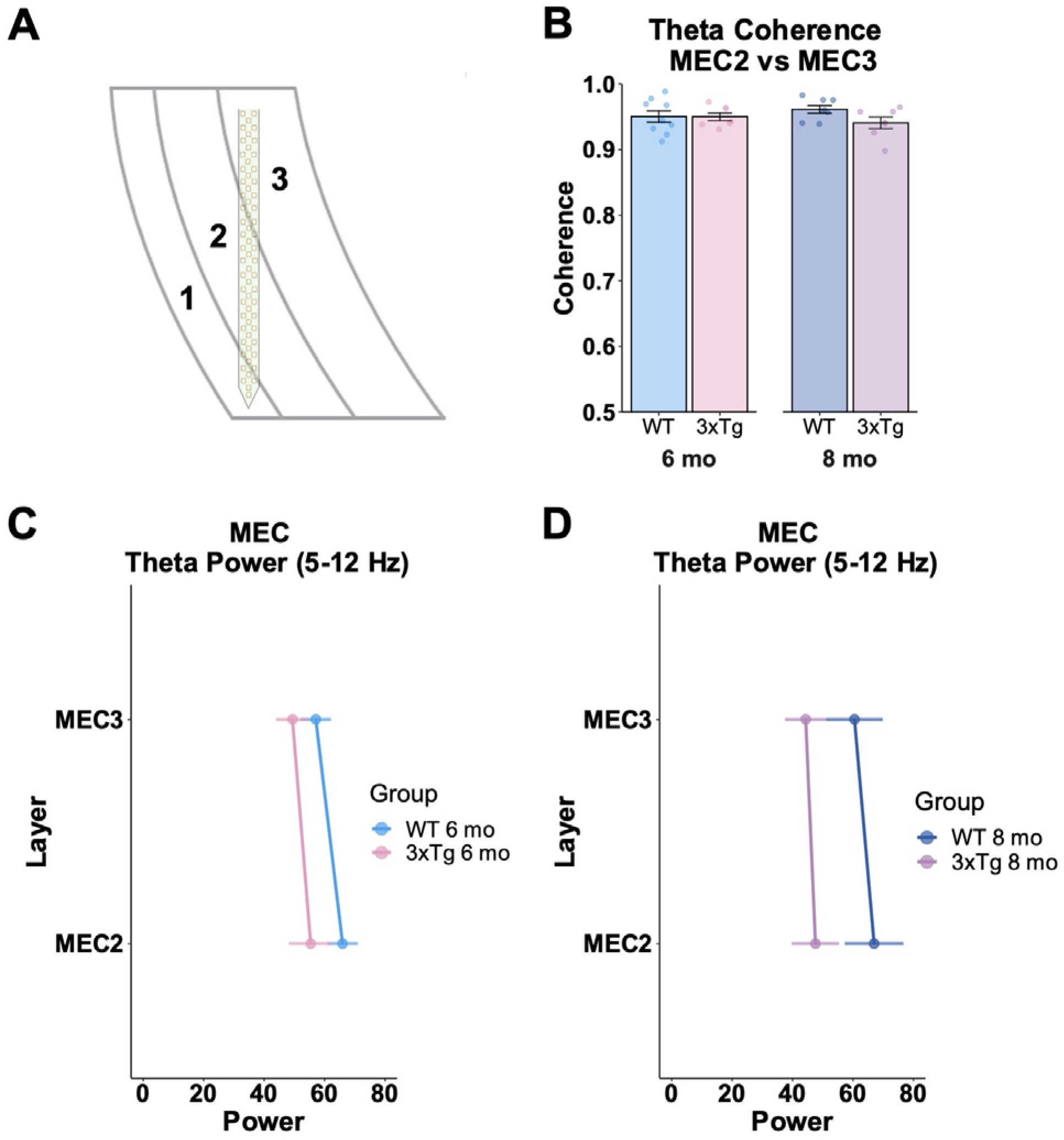
MEC theta power and coherence. **A.** Schematic of probe location in MEC, spanning layers 2 and 3. **B.** Coherence between theta oscillations in MEC2 and MEC3. No changes in theta coherence within MEC across groups. **C.** MEC power by layer in 6-month-old WT and 3xTg mice. No significant genotype effects. **D.** MEC power by layer in 8-month-old WT and 3xTg mice. No significant genotype effects.

**Figure S4 [Related to Figure 3]:**
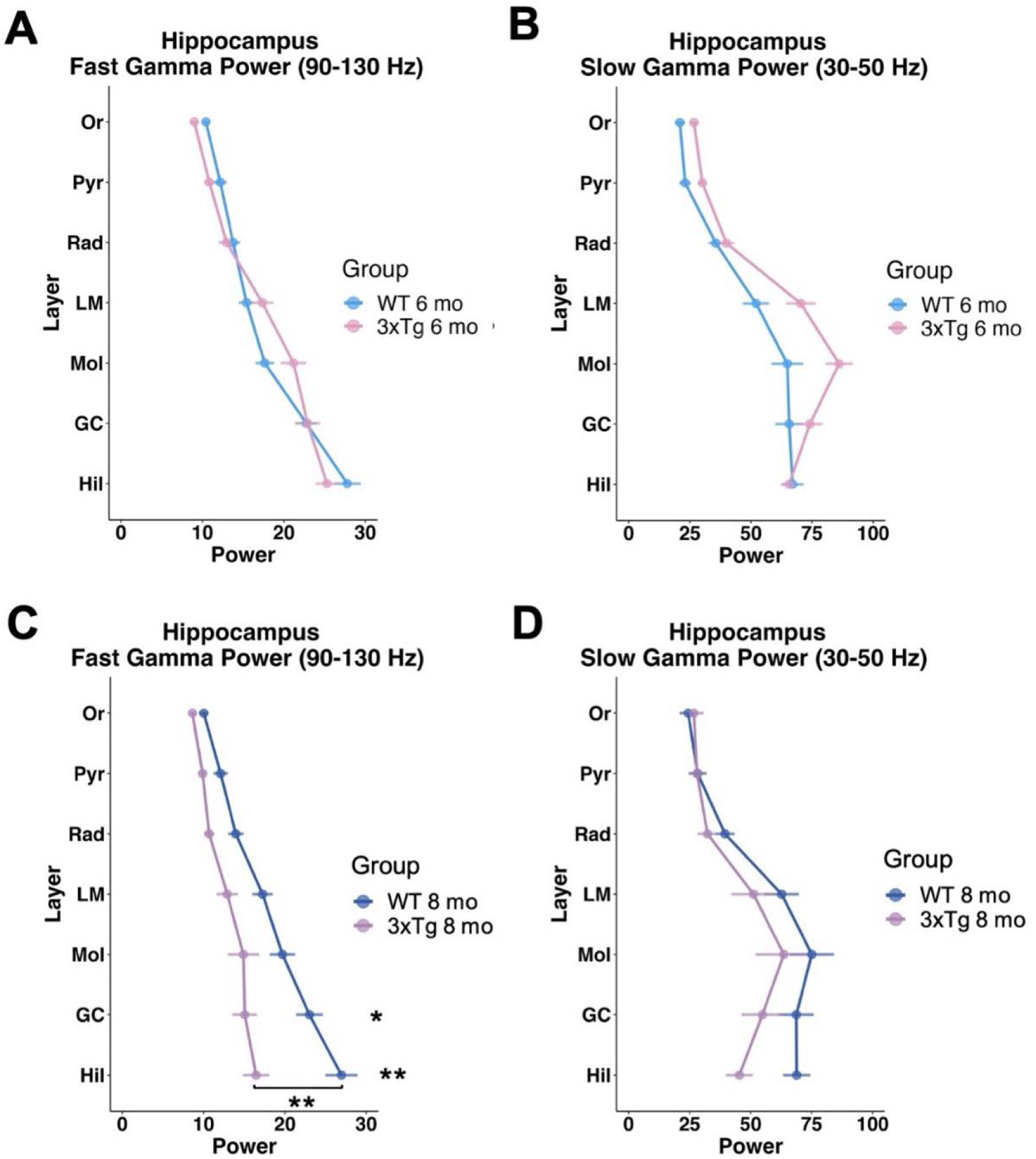
Decreased fast gamma but not slow gamma power in 8-month-old 3xTg mice. **A.** Fast gamma (90-130Hz) power across hippocampal layers in 6-month-old WT and 3xTg mice (two-way ANOVA, layer x genotype interaction F(6, 90) = 6.04, p<0.0001, posthocs by layer corrected with Holm method, all p>0.1). **B.** Slow gamma (30-50 Hz) power across hippocampal layers in 6-month-old WT and 3xTg mice (two-way ANOVA, layer x genotype interaction F(6, 90) = 4.54, p=0.0005, posthocs corrected with Holm method, all p>0.1). **C.** Fast gamma power across hippocampal layers in 8-month-old WT and 3xTg mice. Reduced fast gamma power in 8-month-old 3xTg mice, particularly in granule cell layer (GC) and hilus (Hil), compared to age-matched WT controls (two-way ANOVA, main effect of genotype F(1,15) = 9.43, p = 0.008, layer x genotype interaction F(6, 90) = 7.85, p<0.0001, posthocs corrected for multiple comparisons with Holm method. GC p = 0.017, Hil p = 0.007, Rad p = 0.061, all other layers p>0.1). **D.** Slow gamma (30-50 Hz) power across hippocampal layers in 8-month-old WT and 3xTg mice (two-way ANOVA, layer x genotype interaction F(6,90) = 3.64, p=0.003, posthocs corrected with Holm method, Hil p = 0.067, all other layers p>0.1). Error bars show s.e.m. *p<0.05, **p<0.01.

**Figure S5 [Related to Figure 3].**
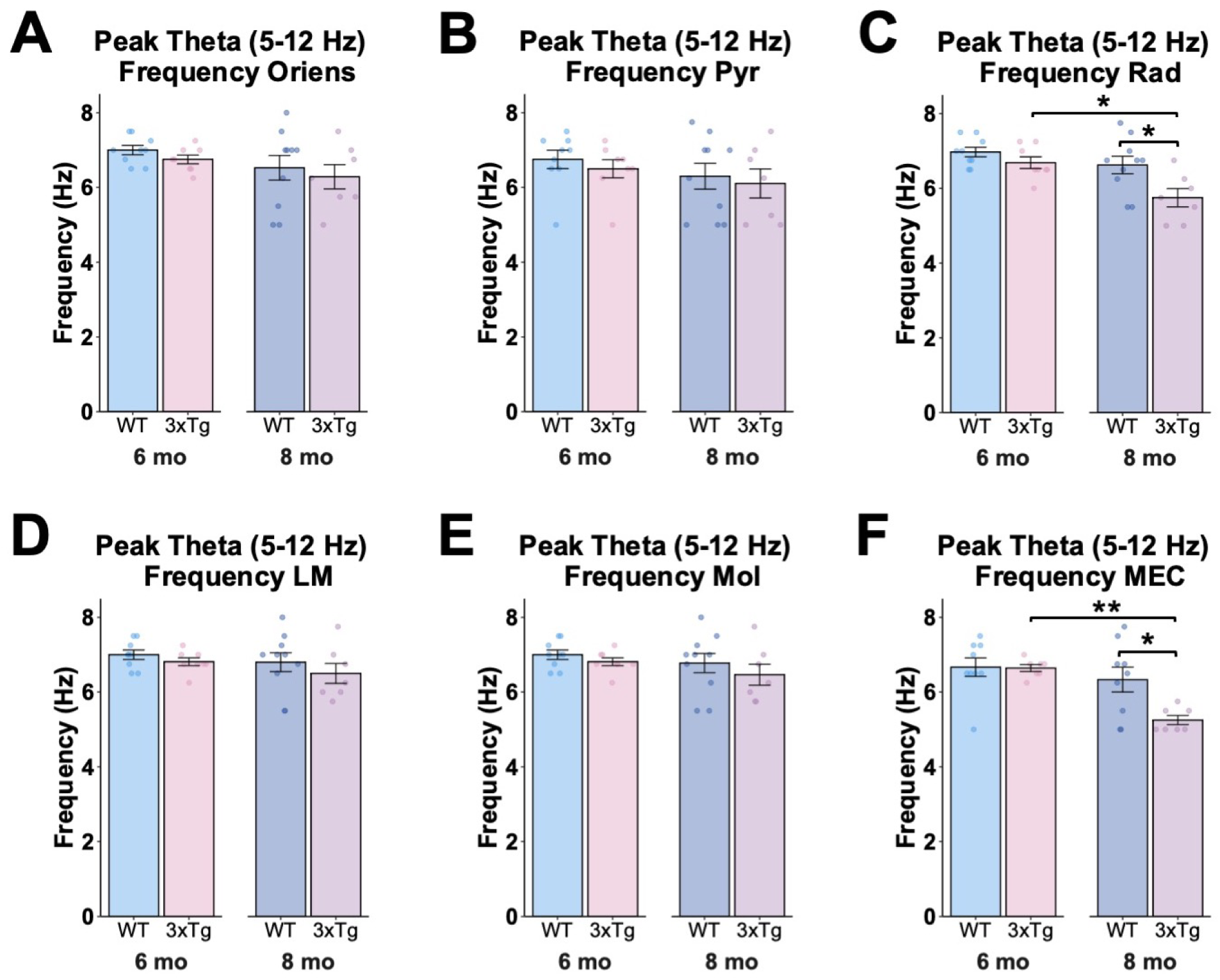
Theta frequency in Hippocampus and MEC. **A.** Peak theta frequency between 5 and 12 Hz by group in CA1 oriens. No significant differences between groups. **B.** Peak theta frequency between 5 and 12 Hz by group in CA1 pyramidal layer. No significant differences between groups. **C.** Peak theta frequency between 5 and 12 Hz by group in CA1 radiatum. Decreased theta-frequency in 8-month-old 3xTg mice (Two-way ANOVA, main effect of age F(1,30) = 10.41, p = 0.003, main effect of genotype F(1,30) = 8.48, p = 0.007; posthocs corrected with Holm method, 3xTg 6 mo vs. 3xTg 8 mo p = 0.015, WT 8 mo vs. 3xTg 8 mo p = 0.015). **D.** Peak theta frequency between 5 and 12 Hz by group in CA1 lacunosum moleculare. No significant differences between groups. **E.** Peak theta frequency between 5 and 12 Hz by group in DG molecular layer. No significant differences between groups. **F.** Peak theta frequency between 5 and 12 Hz by group in MEC. Decreased theta-frequency in 8-month-old 3xTg mice (Two-way ANOVA, age x genotype interaction F(1,28) = 4.63, p = 0.040, posthocs corrected with Holm method, 3xTg 6 mo vs. 3xTg 8 mo p = 0.003, WT 8 mo vs. 3xTg 8 mo p = 0.013). Error bars show s.e.m. *p<0.05, **p<0.01.

**Figure S6 [Related to Figure 4-6].**
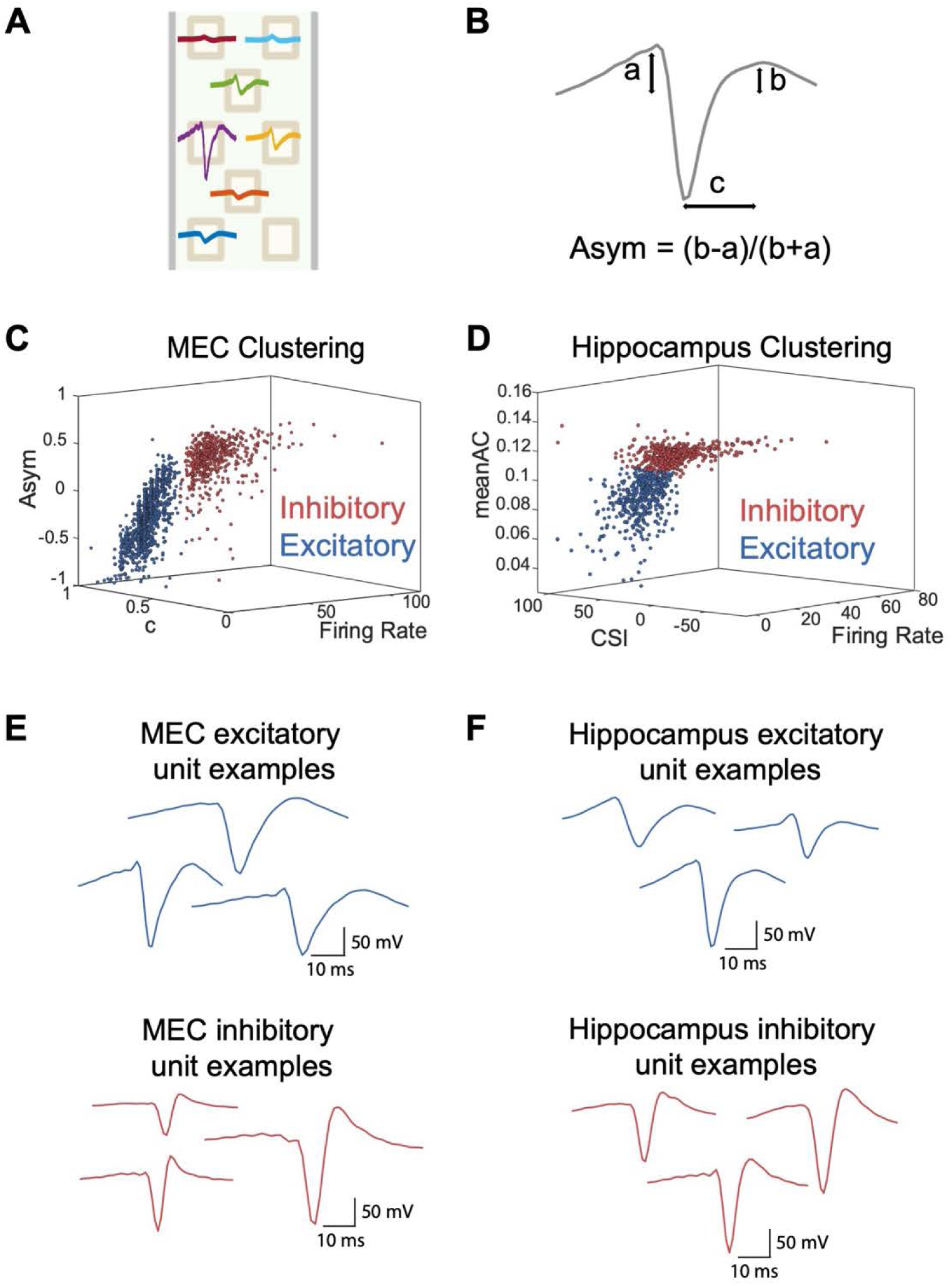
Clustering of excitatory and inhibitory neurons. **A.** Example waveforms of a putative neuron recorded across seven silicon probe channels and clustered using Kilosort 2.5. **B.** Schematic depicting trough to peak latency (c) and waveform asymmetry (asym). **C.** Clustering of MEC units into putative excitatory (blue) and inhibitory (red) units. Clusters were determined using k-means clustering based on trough to peak latency of waveforms (c). **D.** Clustering of hippocampal units into putative excitatory (blue) and inhibitory (red) units. 3D plot shows values for mean firing rate (FR), mean autocorrelation (meanAC), and complex spike index (CSI) for each unit. Clustering was performed using thresholds for FR, meanAC, CSI, and trough to peak latency of waveforms (c). **E.** Example waveforms from MEC excitatory and inhibitory units**. F.** Example waveforms from hippocampal excitatory and inhibitory units.

**Figure S7 [Related to Figure 4-6].**
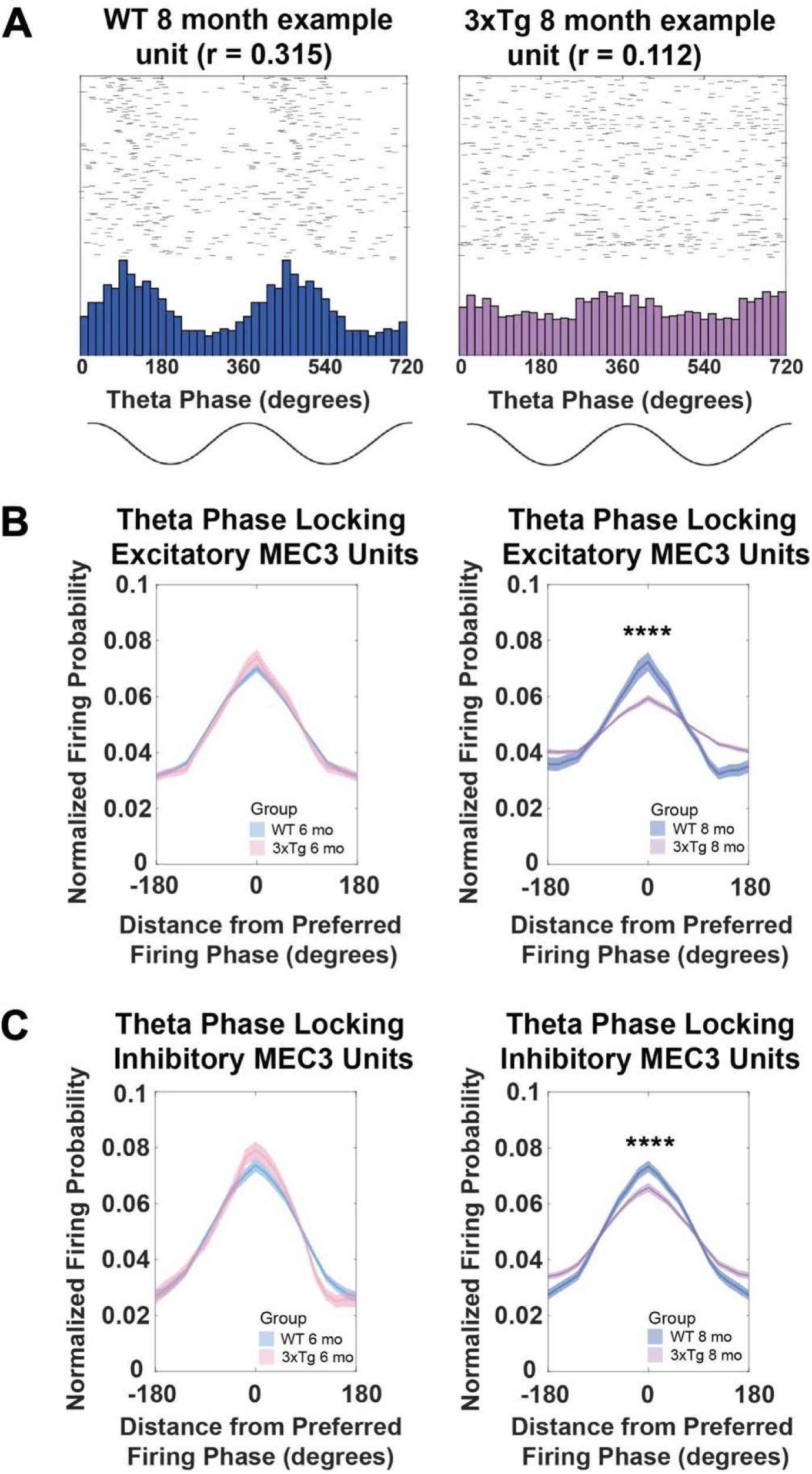
Additional visualizations of theta phase locking A. Phase locking of example units from a WT animal (left) and 3xTg animal (right). Histograms show normalized probability of firing at a given theta phase. Raster plots show spike times across 200 theta cycles in which the unit fired. Data is double plotted to aid in visualizing theta phases. **B.** Normalized firing probability averaged across all excitatory MEC3 units in each genotype at 6 months (left) and 8 months (right), where theta phase was divided into 21 equal bins and each unit’s distribution was centered around its most preferred theta phase and smoothed prior to averaging. Note the flatter distribution in 8 mo 3xTg units, reflecting a weaker phase preference. (WT 8 mo vs. 3xTg 8 mo: Two-way ANOVA, genotype x phase interaction F(20, 2120) = 12.01, p < 0.0001). **C.** Same as B for inhibitory MEC3 units (WT 8 mo vs. 3xTg 8 mo: Two-way ANOVA, genotype x phase interaction F(20,2000) = 6.07, p < 0.0001). Error bars show s.e.m. ****p<0.0001.

**Figure S8 [Related to Figure 4-5].**
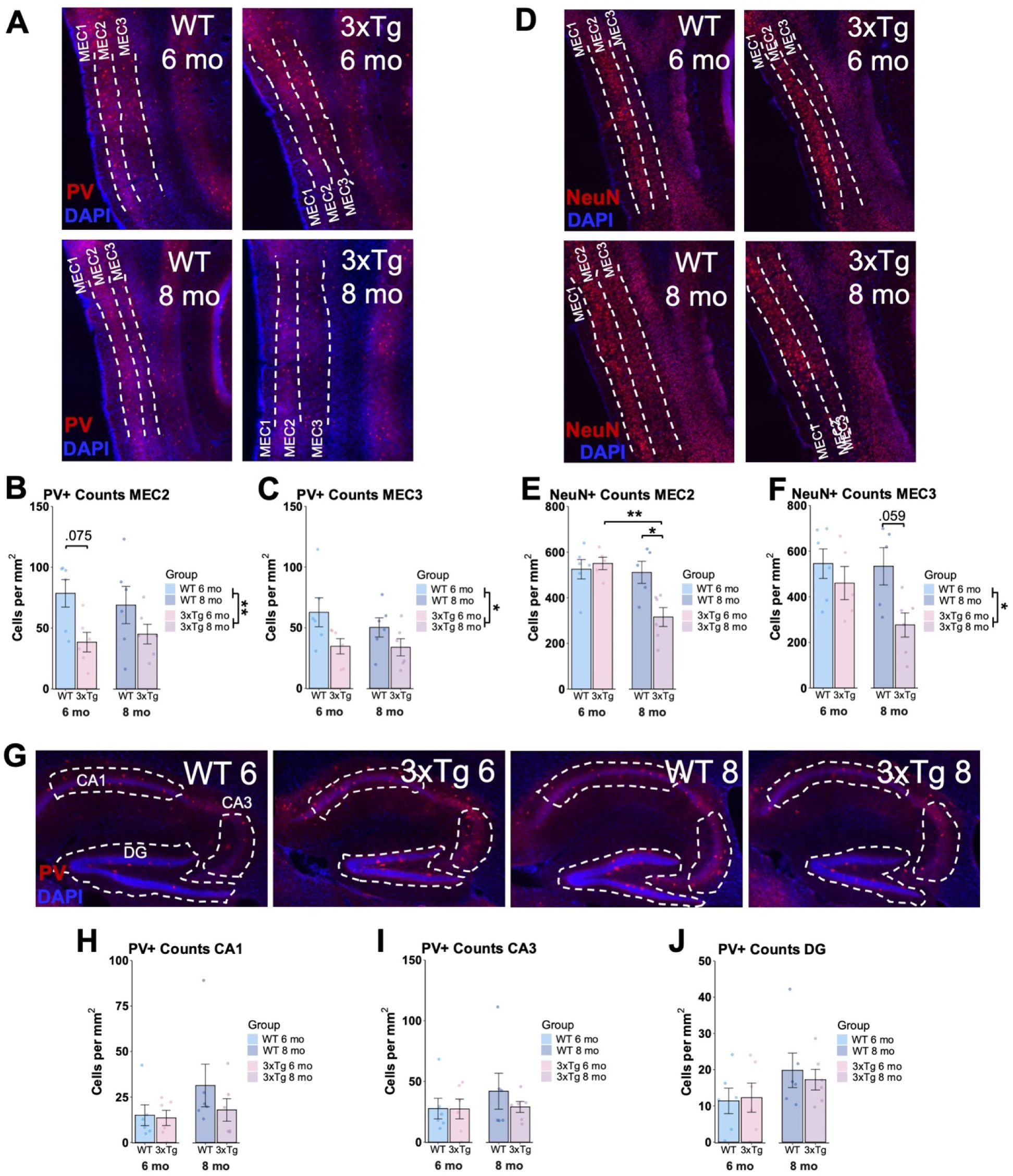
Decreased PV+ interneuron counts in MEC of 3xTg mice as early as 6-months, but no changes in hippocampal PV+ counts. **A.** Example representative PV immunohistochemistry images from MEC of 6 (top) and 8 (bottom)-month-old WT (left) and 3xTg (right) mice. Blue = DAPI, Red = Parvalbumin (PV). **B.** PV+ cell counts in MEC2 in 6 and 8-month-old 3xTg and WT mice. Counts were reduced in 3xTg mice (Two-way ANOVA, main effect of genotype F(1,20) = 8.34, p= 0.009, posthocs corrected with Holm method WT 6 mo vs. 3xTg 6 mo p = 0.075). **C.** PV+ cell counts in MEC3 were reduced in 3xTg mice (Two-way ANOVA, main effect of genotype F(1,20) = 6.71, p = 0.017). **D.** Example representative NeuN immunohistochemistry images from MEC of 6 (top) and 8 (bottom)-month-old WT (left) and 3xTg (right) mice. Blue = DAPI, Red = NeuN. **E.** NeuN+ cell counts in MEC2. Decreased neuronal counts in 8-month-old 3xTg mice (Two-way ANOVA, age x genotype interaction F(1,18) = 7.22, p = 0.015, posthocs corrected with Holm method, 3xTg 6 mo vs. 3xTg 8 mo p = 0.003, WT 8 mo vs. 3xTg mo p = 0.010). **F.** NeuN+ cell counts in MEC3 (Two-way ANOVA, main effect of genotype F(1,18) = 6.44, p = 0.021, posthocs corrected with Holm method, WT 8 mo vs. 3xTg 8 mo p = 0.059). **G.** Representative PV immunohistochemistry images from hippocampus. **H.** PV+ cell counts in CA1. **I.** PV+ cell counts in CA3. **J.** PV+ cell counts in DG. No differences between groups in hippocampus. **Sample sizes:** MEC and Hippocampus PV: N=6 per group, 3 males and 3 females. MEC NeuN: WT 6 mo (N = 6, 3M/3F), WT 8 mo (N= 5, 3M/2F), 3xTg 6 mo (N = 5, 3M/2F), 3xTg 8 mo (N=6, 3M, 3F).

**Figure S9 [Related to Figure 6].**
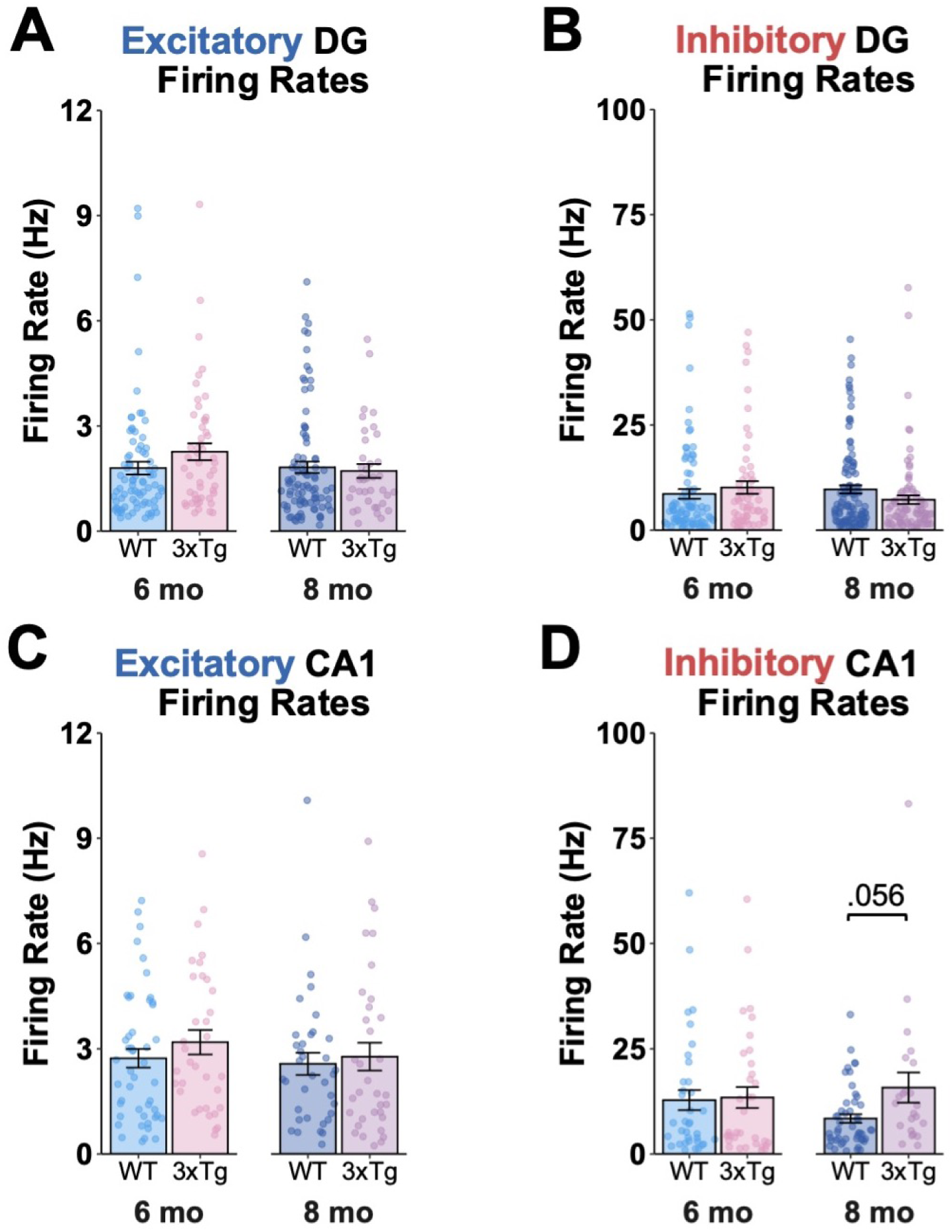
No changes in hippocampal firing rates. **A.** Firing rates by group for excitatory DG units. **B.** Firing rates by group for inhibitory DG units. **C.** Firing rates by group for excitatory CA1 units. **D.** Firing rates by group for inhibitory CA1 units. (Mann-Whitney test, WT 8 mo vs. 3xTg 8 mo adj p = 0.056). All other comparisons p>0.05 after controlling for multiple comparisons.

## Notes

### Competing Interest Statement

The authors have declared no competing interest.

